# DexDesign: A new OSPREY-based algorithm for designing *de novo* D-peptide inhibitors

**DOI:** 10.1101/2024.02.12.579944

**Authors:** Nathan Guerin, Henry Childs, Pei Zhou, Bruce R. Donald

## Abstract

With over 270 unique occurrences in the human genome, peptide-recognizing PDZ domains play a central role in modulating polarization, signaling, and trafficking pathways. Mutations in PDZ domains lead to diseases such as cancer and cystic fibrosis, making PDZ domains attractive targets for therapeutic intervention. D-peptide inhibitors offer unique advantages as therapeutics, including increased metabolic stability and low immunogenicity. Here, we introduce DexDesign, a novel OSPREY-based algorithm for computationally designing *de novo* D-peptide inhibitors. DexDesign leverages three novel techniques that are broadly applicable to computational protein design: the Minimum Flexible Set, K*-based Mutational Scan, and Inverse Alanine Scan, which enable exponential reductions in the size of the peptide sequence search space. We apply these techniques and DexDesign to generate novel D-peptide inhibitors of two biomedically important PDZ domain targets: CAL and MAST2. We introduce a new framework for analyzing *de novo* peptides—evaluation along a replication/restitution axis—and apply it to the DexDesign-generated D-peptides. Notably, the peptides we generated are predicted to bind their targets tighter than their targets’ endogenous ligands, validating the peptides’ potential as lead therapeutic candidates. We provide an implementation of DexDesign in the free and open source computational protein design software OSPREY.

## 1 Background and Introduction

Since the 1921 discovery^1^ of the peptide hormone insulin to treat diabetes, many peptides and peptide-derived therapeutics have come into clinical use, with more than 30 achieving final regulatory approval since the year 2000^2^. The use of peptides as therapeutics has a number of advantages, including standard protocols for synthesis, good efficacy, high potency, and selectivity^3,4^. On the other hand, peptide therapeutics have a number of drawbacks, including poor stability, oral bioavailability, membrane permeability, and retention^4^. Changing the chirality of L-amino acids to their D counterparts in peptides is a strategy medicinal chemists have used to address these shortcomings.

In Section 1.1, we describe the benefits of incorporating D-amino acids into therapeutic peptides. Section 1.2 provides background on PDZ domains that researchers have investigated targeting with L-peptides for biomedical purposes. Section 1.3 describes previous computational protein redesign software and algorithms for designing proteins and peptides incorporating non-canonical and D-amino acids. Section 0 concludes with a summary of DexDesign, a new algorithm we developed and incorporated into the protein design software OSPREY. With the necessary background covered, the rest of this article focuses on an application of the DexDesign algorithm to generate *de novo* D-peptide inhibitors of two biomedically important PDZ domains targets: CAL and MAST2. We then evaluate each computationally generated peptide using multiple structural criteria, including predicted binding affinity and whether a D-peptide mimics binding interactions previously shown to be important to L-peptide binding to PDZ domains.

### 1.1 Benefits of including D-amino acids in peptides

The inclusion of D-amino acids can increase peptide stability by decreasing the substrate recognition by proteolytic enzymes^5^. For example, Chen et al. improved both stability and binding affinity of a bicyclic peptide inhibitor of the cancer-related protease urokinase-type plasminogen activator by substituting a single D-serine for a glycine^6,7^. Haugaard-Kedström et al. observed that the simple substitution of D-amino acids in two positions of their *de novo* PDZ domain inhibitor greatly improved metabolic stability by increasing their peptide’s half-life 24-fold^8^. More ambitiously, Liu et al. constructed an entirely D-peptide inhibitor of the MDM2 oncoprotein using mirror image phage display, an experimental technique used to discover D-peptide drug candidates, that inhibited growth of glioblastoma both in cell culture and nude mouse xenograph models^9^. Nevertheless, the challenge of preparing an enantiomeric protein target for mirror image phage display remains a drug-discovery bottleneck^10^.

### 1.2 PDZ domains

With over 270 unique occurrences in more than 150 human proteins, PDZ domains constitute the largest family of peptide-recognition domains in the human genome^11^. A typical PDZ domain has 80-100 amino acids folded into five core β-strands (β1-β5) and two α-helices (α1 and α2)^11–13^. They facilitate a variety of cellular functions, such as modulating polarization, signaling, and trafficking pathways through interaction with short linear motifs (SLiMs) located at the C-terminus of their ligands^11,14–16^. Usually, SLiMs bind into a groove of the PDZ domain between α2 and β2, extending the β2/β3 sheet^14^. Modulating the interaction between a SLiM and its PDZ binding partner is a strategy that both viruses and therapeutics aim to exploit^11,17^ and has been explored by previous computational design techniques^18–26^.

### 1.2.1 CFTR-associated ligand

Cystic fibrosis can cause serious pulmonary and respiratory problems in the lungs by causing the development of a thick mucus that promotes bacterial infection and inflammation. It is caused by many mutations in the cystic fibrosis transmembrane conductance regulator (CFTR), such as ΔF508, which causes destabilized, misfolded CFTR^18,27^. The CFTR-associated ligand (CAL) binds CFTR via CAL’s PDZ domain (CALP), which shepherds CFTR through rapid degradation via a lysosomal pathway^18^. A number of research groups have developed peptide stabilizers that bind to CALP, preventing CFTR lysosomal trafficking and degradation. Our lab used computational peptide design to develop a hexamer that bound 170-fold more tightly to CALP than CALP bound the CFTR C-terminus, rescuing CFTR activity in monolayers of polarized human upper airway epithelial cells that contain the ΔF508 deletion in CFTR—80% of cystic fibrosis patients are homozygous for this mutation^18,28^. Cyclic peptides targeting CALP have also been developed—Dougherty et al.^29^ developed a highly selective stabilizing cyclic peptide that binds CALP with a K_D_ of 6 nM.

### 1.2.2 MAST2

During viral infection, the rabies virus exploits SLiM/PDZ-domain interactions to further its propogation^30,31^. In a neuron, phosphatase and tensin homolog deleted on chromosome 10 (PTEN)’s SLiM interacts with the PDZ domain of microtubule-associated serine-threonine kinase 2 (MAST2) to regulate pathways inhibiting neuronal survival, regrowth, and regeneration^32,33^. The rabies virus glycoprotein’s C-terminal residues interact with MAST2’s PDZ domain, disrupting the ability of MAST2 and PTEN to form a complex and inhibit neurite outgrowth and apoptosis^30,33,34^. Recognizing the therapeutic potential of promoting neurite outgrowth in the treatment of neurodegenerative disease, Khan et al. developed three peptides that mimic and improve upon the rabies virus glycoproteins’ interaction with MAST2’s PDZ domain, stimulating neurite outgrowth in proportion to the affinity the peptide bound MAST2^33^.

### 1.3 Computational tools and algorithms for designing D-peptides

OSPREY is a free and open-source software program containing a suite of computational protein design algorithms developed in our lab^35^. OSPREY has been used to, among other things, predict resistance mutations ablating efficacy of antibiotics used to treat methicillin-resistant *Staphylococcus Aureus*^36,37^ and small molecule inhibitors used to treat melanoma, lung, stomach and colorectal cancers^38,39^, design and structurally characterize peptide inhibitors of CALP for treating CFTR^18,28^, and improve broadly neutralizing antibodies against HIV-1^40,41^. OSPREY has been used^42–44^ to computationally redesign proteins with canonical and non-canonical amino acids, as well as to optimize protein:small molecule interactions^38,39,45^, but until now has not had the capability to design D-peptides. Given the promising biomedical potential of D-peptides^9,46–50^, extending OSPREY’s ensemble-based, provable protein design algorithms to D-peptide design could greatly decrease the required quantity of expensive, time-intensive experiments.

In addition to the expensive, purely experimental techniques for D-peptide design, such as mirror phage display reviewed in Sec. 1.1, there are a few previous computational techniques for D-peptide design (also reviewed in Donald 2011, Chapter 9)^51^. Noncanonical amino acids have been implemented in protein design algorithms, such as OSPREY, as early as 2006^52^. Elkin et al used the Multiple Copy Simultaneous Search^53^ method to predict candidate D-peptide inhibitors of hepatitis delta antigen dimerization^54^. Recent versions of Rosetta have included functionality to incorporate non-canonical and D-amino acids^55,56^. Philip Kim’s group developed a computational D-peptide design technique based on creating a mirror image of the PDB, identifying hotspot interactions, and searching the D-PDB for similar configurations of hotspot residues^57,58^. Overall, the number of algorithms the protein designer has available for D-peptide design is notably sparser than for L-design, and the development of additional computational protocols for this important task is warranted.

### 1.4 DexDesign

In this paper we present an algorithm, DexDesign, for designing *de novo* D-peptides in OSPREY (Figure 1). DexDesign constructs D-peptide scaffolds by mirroring the structure of a L-protein:peptide complex into D-space, then uses the geometric search algorithms in MASTER^59^ to search hundreds of thousands of L-protein structures for substructures with backbones similar to the D-peptide. It then uses the iMinDEE/K* algorithm^60,61^, in OSPREY to redesign a scaffold D-peptide’s sidechains to optimize target binding. Given the biomedical importance of modulating CALP and MAST2 PDZ domain interactions, coupled with the advantages of D-peptide therapeutics, we use DexDesign to predict D-peptides inhibitors of these two protein targets.

**Figure 1:**
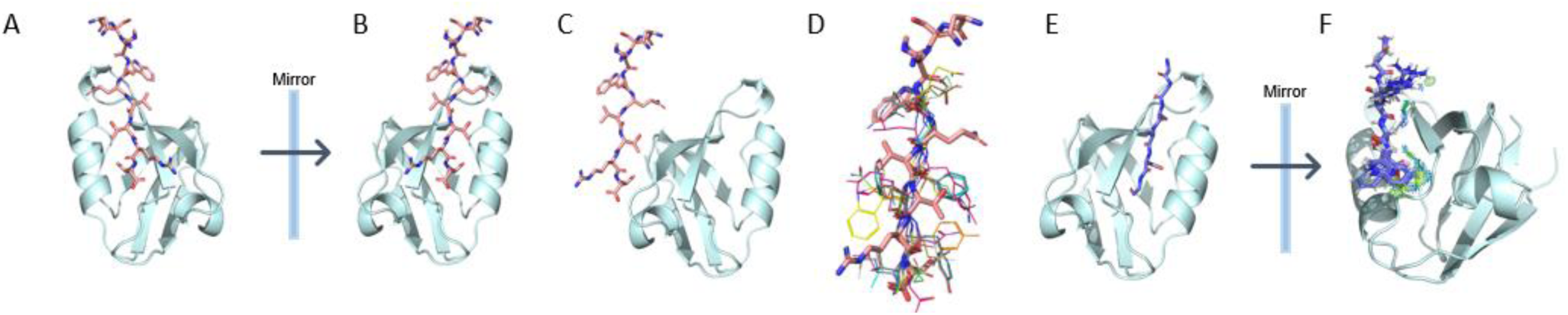
Example of the DexDesign protocol applied to CALP. **a)** The potent inhibitor kCAL01 (in pink) bound to CALP (in cyan) (PDB ID 6ov7) was used as a starting point for a DexDesign search for a D-peptide inhibitor^28^ of CALP. Both kCAL01 and CALP are composed of solely L-amino acids. **b)** The input structure is reflected to produce a mirror-image of the kCAL01:CALP complex, which flips the chirality of all amino acids to D. **c)** The complex is split into its constituent peptide and protein components. **d)** Residues P^0^ to P^-5^, which are the residues located within CALP’s binding pocket, are used as the query structure to conduct a MASTER^60^ search of a large database of L-protein structure to find substructures with similar backbones (as determined by backbone RMSD) to the D-version of kCAL01. 10 representative matches of L-peptide segments (in multicolored wire representation) are overlaid on the pink stick D-version of kCAL01. **e)** Each L-peptide match is aligned to the D-version of kCAL01 in the D-kCAL01:CALP complex structure and D-kCAL01 is removed. Shown (in purple sticks) is an L-peptide substructure GGAASG (residues 168-173) that MASTER identified in Mycobacterium tuberculosis Rv0098 (PDB ID 2PFC). This L-peptide^59^ forms the basis for OSPREY redesign. **f)** The L-peptide:D-CALP complex is reflected once again to form a D-peptide:L-CALP complex. The K* algorithm^60,61^ in OSPREY^34^ is then invoked to conduct a search over D-peptide sequences and continuous sidechain conformations to optimize the D-peptide for binding. K* identified two mutations at positions P^0^ and P^-2^ predicted to improve binding of the peptide with a normalized ΔΔG of -1.4 kcal/mol, improving KD by 9-fold (see Appendix B for information on the normalization procedure). Position P^0^ is mutated from Gly to Trp, and P^-2^ is mutated from Ala to Arg. Shown is an OSPREY-predicted low energy ensemble of the D-peptide GGARSW with Molprobity probe dots^62–64^ showing goodness-of-fit the OSPREY-predicted mutated D-sidechains make with CALP.

## 2 Methods

### 2.1 Algorithm and Computational Protocol

DexDesign generates *de novo* D-peptides by combining MASTER’s molecular structure search^59^ with novel improvements (described herein) to provable computational protein redesign algorithms in OSPREY^35,51^, mediated via an energy-equivariant geometric transformation (EEGT). EEGTs, including translation, rotation, and reflection, are isometries of a molecular structure that do not change the energy of the structure. Each EEGT corresponds to a symmetry in the energy field^62^. For example, an energy function computes the same energy of protein structure *s* and *s* reflected over the Cartesian *x-y* plane. Now, the MASTER algorithm searches a database of protein structures for a user-specified query structure and is guaranteed to find all protein substructures in the database with a backbone RMSD below a cutoff threshold^59^. The K* algorithm in the OSPREY software suite^35^ searches for amino acid substitutions that maximize a design objective, such as binding affinity or specificity^60,63–65^. It does this by exploiting molecular ensembles to compute a provably accurate ε-approximation to the model’s theoretical binding constant, 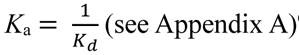^60,63^.

In essence, DexDesign invokes MASTER search as a subroutine to generate D-peptide scaffolds with backbone conformations similar to their L-peptide counterpart, then invokes K* as a subroutine to optimize amino acid sequences and side chain conformations on those scaffolds. After preparing a database (DB) of L-protein structures, a protein designer initiates DexDesign by identifying a protein target (*t*) of interest, for which there exists a structure of a protein or peptide (*p*) bound to *t*. In MASTER terminology, the structure of this bound complex will become our query, *q_tp_*. Below, we define terms for the DexDesign algorithm specified in Figure 2:

**Figure 2:**
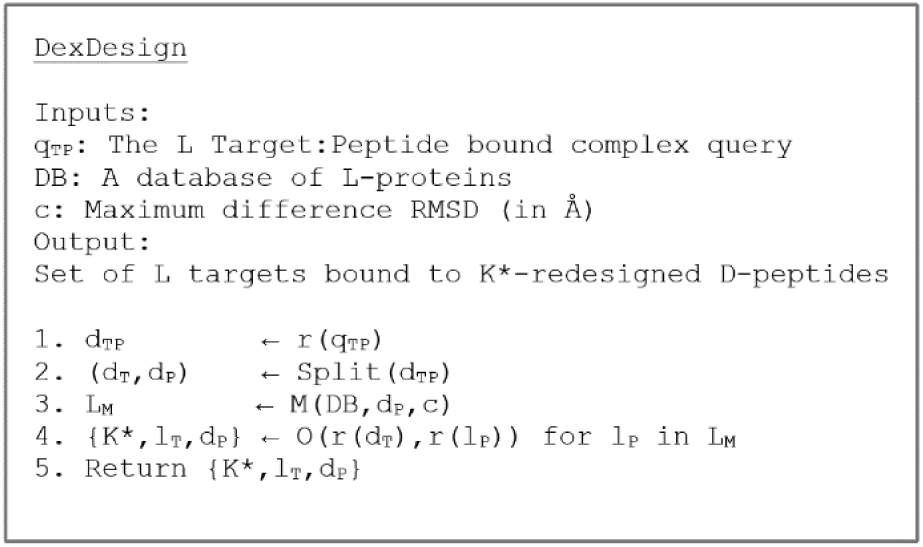
The DexDesign Algorithm. DexDesign takes as input a query structure (q) of a bound target (t) and peptide (p), a database (DB) of L-protein structures, and a cutoff (c). Line 1 reflects the L-protein complex into D-space. Line 2 splits the target and peptide into two structures, the D-target (dT) and the D-peptide (dP). Line 3 calls MASTER^65^ (M) to search the L-protein database for all substructures with a backbone RMSD to dP less than c. The set of results is saved in LM. Line 4 reflects each MASTER peptide (lP) and D-target (dT), to make a D-version of lP bound to the original L-target. OSPREY^34^ K* redesign^41,66^ is then run on each target:peptide complex (lT + dP), resulting in a set of K* scores (K*), along with an OSPREY-predicted structural ensemble of the D-peptide (dP) and L-target (lT) complexes, with the sequence and continuously minimized sidechains of dP optimized to bind lT. The K* scores and computed structural ensembles are returned on Line 5. While this novel control flow is surprisingly simple, a key contribution of this paper is finding subroutines to make the steps both efficient and provably accurate, properties that are rare in computational structural biology.

1. Let *S*_*n*_ be a protein structure with *n* residues. We define substructure *S*_*i,j*_ of *S*, where 1 ≤ *i* < *j* ≤ *n*, to be a structure of residues *i* through *j* of *s*.
2. Let *r*(*S*, *a*) be a function that reflects all atoms in protein structure *S* across a plane *a*. Without loss of generality, we let *a* be the *x-y* plane and define *r*(*S*) = *r*(*S*, *a*) henceforth. Then, *r*(*S*)reflects all L-amino acids, including those with multiple stereocenters (*viz.*, threonine and isoleucine), into their corresponding D-versions, and vice-versa (see Appendix Figure 1 for an example).
3. Let *M*(*DB*, *s*, *c*) be the MASTER subroutine. *M* returns a set of substructures from DB with backbone RMSD, when optimally aligned with protein substructure *s,* less than *c* Å.
4. Let *O*(*p, t*) be the OSPREY K* subroutine. *O* redesigns peptide *p* towards increased binding affinity with protein target *t* by searching over mutated and continuously minimized amino acid sidechains, and returns a set of mutant sequences (and structural molecular ensembles) derived from *p* that have improved binding with *t*.

### 2.2 New features in OSPREY: Customizable conformation libraries and D-protein/peptide design

In previous work^38,39,66–68^, a typical OSPREY-based computational protein redesign entailed: 1) selecting a starting molecular structure, 2) adding hydrogens, 3) specifying the design algorithm and its input parameters, 4) running the algorithm, and 5) analyzing the results. To enable Step 4, OSPREY included a default library of amino acid atom connectivity templates from Amber^69^ and rotamers from Lovell et al^70^. These templates and rotamers then became starting points for continuous minimization within a voxel during the sequence and computational search^60,71^. Embedding the templates within the algorithm provided protein designers with simple defaults for the majority of protein redesign problems. And while some works^41,72,73^ have expanded these defaults in certain cases, such as in our use of non-canonical amino acids to design CALP inhibitors^72^, the necessity of providing a simple, general approach that enabled protein designers to experiment with diverse and novel biochemical building blocks remained. The implementation of a general, in contrast to application-specific, approach to modeling templates and flexibility enables designers to design proteins with chemistries that the creators of the protein design software didn’t even anticipate!

To meet this need, we have simplified the process of specifying rotamers, voxel-based continuous minimization, new molecular fragment templates, and even entire conformation libraries in OSPREY. OSPREY continues to provide intelligent defaults, but they are moved from deep within the software and are now exposed to the designer, allowing the designer to modify them as needed in a simple graphical user interface (Appendix Figure 2, right). This seemingly simple change has profound implications for OSPREY. When the complete conformation space specification (i.e., the design parameters such as the mutable residues, the flexible residues, etc.) is a user-modifiable input to the algorithm, new classes of design capabilities, such as design with D-amino acids via DexDesign, are unlocked.

The molecular interaction forces between two molecules are invariant over a reflection of those two molecules. Put another way, if *K*_*D*_(*x*, *y*) is the dissociation constant for protein *x* binding protein *y,* then *K*_*D*_(*x*, *y*) = *K*_*D*_(*r*(*x*), *r*(*y*)). The OSPREY energy function, as described in detail in previous works,^35,61,74,75^ mimics this physics precisely be being (exactly) energy equivariant with respect to reflection, allowing us to add the ability to design D-proteins and peptides in OSPREY. We accomplished this by reflecting OSPREY’s default L-conformation library into D-space. A protein designer can now use the functionality described in Section 2.2.1 to specify a D- or L-conformation library on a per-protein basis in a single design (Appendix Figure 2, left).

### 2.3 Applying DexDesign to CALP and MAST2

To use DexDesign to predict *de novo* D-peptide inhibitors to CALP and MAST2, we started with structures of two complexes: kCAL01 bound to CALP (PDB ID 6ov7)^28^ and PTEN bound to MAST2 (PDB ID 2kyl).^76^ We created a database of high-resolution L-protein structures by mining the RCSB PDB^77^ for crystallographically-determined structures with a resolution better than 2.5 Å, omitting DNA, RNA, and small molecules. This resulted in a database containing 119,160 structures (see Appendix Figure 6 for further details).

Using the DexDesign algorithm in Figure 2, we first reflected each molecular structure to D-space and split the peptide and target PDZ domain into two separate structures, *d_p_* and *d_t_*, respectively. We then used the MASTER algorithm^59^ to query the database for L-protein substructures with backbones similar to *d_p_*. MASTER returns a set of candidate L-peptides (L_M_), each of which (*l_p_* ∈ L_M_) we superimposed over *d_p_* in the *d_p_*:*d_t_* complex and subsequently removed *d_p_*. We then again reflect each bound complex *l_p_*:*d_t_*, resulting in a *de novo* D-peptide candidate *r*(*l*_*p*_) bound to the original L-protein target *r*(*d*_*t*_) in a complex *r*(*l*_*p*_):*r*(*d*_*t*_).

Prior to executing Step 4 of the DexDesign algorithm (K* redesign; see Figure 2) we further pruned the set of D-peptide candidates based on two additional criteria. First, we visualized candidate D-peptide:L-protein complex structures in PyMol^78^ using Molprobity dots^79,80^ in our lab’s Protein Design Plugin^81^ to evaluate the number and severity of steric clashes, as steric clashes need to be resolved via sequence mutation and additional modeling of continuous side chain flexibility in the K* algorithm. Since clash resolution and peptide improvement via K* redesign utilize the same algorithmic technique and therefore draw from the same pool of limited computational resources, we chose to prioritize D-peptide candidates with fewer clashes so we could allocate more computational resources to improving a D-peptide candidate via K* sequence redesign. Second, we observed instances where MASTER found identical D-peptide candidate sequences with nearly identical structures in multiple distinct PDB files, which we resolved by deduplicating the results.

Using the above criteria, we selected eight promising D-peptide candidates to use as starting points for OSPREY K* redesign. We call these selected candidates *D-peptide redesign (DPR) scaffolds*. The complete set of DPR scaffolds are described in Appendix Table 1. To evaluate and improve upon the DPR scaffolds, we developed three new design techniques. Using the series of pruning steps and approximations described herein, we show the DexDesign algorithm implements exponential reductions in the size of peptide search space, obtaining empirical running times with a median of about six minutes.

#### 2.3.1 New Design Techniques: Minimum Flexible Set, Inverse Alanine Scanning, & Mutational Scanning

The following design techniques assume that the protein designer has a fixed amount of time and computational resources at their disposal. To that end, they are formulated to allow designers to rapidly evaluate their DPR scaffolds by restraining the conformation space to the minimal size necessary to computationally evaluate a hypothesis. Conformation space size grows exponentially with the number of flexible residues. Here, restricting the size of a conformation space is an effective technique to obtaining computational predictions quickly. Further, DexDesign’s *de novo* peptide design has one important distinction from protein redesign: the model of the starting protein structure used as input for K* redesign is a theoretical model, rather than one determined by experiment. To address this challenge—which is inherent to *de novo* design—we have developed new design techniques that systematically evaluate the quality of DPR scaffolds and rigorously suggest mutations that are predicted to improve the D-peptide’s binding affinity to its target PDZ domain.

##### 2.3.1.1 Design Technique 1: Identifying a Minimum Flexible Set

The MASTER search (see Figure 2) ranks by backbone-only RMSD. Hence, as visualized in Figure 3 (A), the L-peptide search results often contain side chains in sterically unfavorable positions that clash with the D-protein target. We call the unique set of residues that must be modeled as continuously flexible^60,82^ to resolve clashes from unpruned candidates the *Minimum Flexible Set (MFS)*. Given a fixed budget of computational resources, MFS enables selection of DPR scaffolds requiring smaller Minimum Flexible Sets, because such DPR scaffolds allow K* to expend more compute resources on searching for favorable mutations that increase binding to the target protein. See Figure 3 (A) for an example of specifying the Minimum Flexible Set for a CALP-DPR candidate.

**Figure 3:**
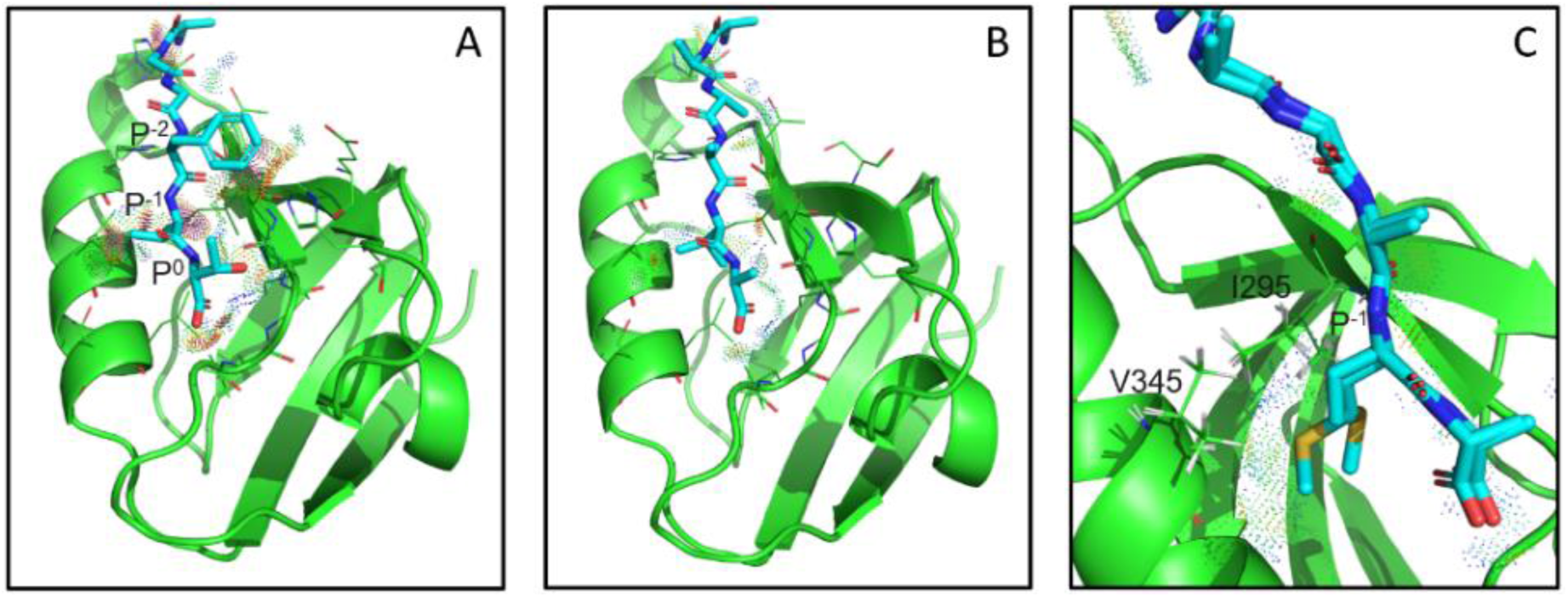
Illustration of the Minimum Flexible Set and Inverse Alanine Scanning design techniques applied to CALP-DPR5. **A:** Determining the Minimum Flexible Set. The CALP-DPR5 scaffold peptide (in cyan) is extracted from a crystal structure of the tobacco necrosis virus (PDB ID 1c8n, residues C61-66, AGGFVT)^81^. When aligned and superimposed over the query peptide kCAL01 and CALP^28^ (in green) to create the DPR scaffold, sidechain and backbone clashes are present that must be resolved (red and pink MolProbity dots)^63,64,80^. The peptide residues that clash with CALP are located at P^0^ (Thr), P^-1^ (Val), P^-2^ (Phe), and P^-5^ (Ala). The *Minimum Flexible Set,* or those residues that must be allowed to undergo continuous minimization (see Gainza et al, 2012)^66^ in all designs derived from CALP-DPR5, are the peptide residues located at P^0^, P^-1^, and P^-2^. The clash involving the alanine located at P^-5^ can be resolved by using OSPREY to translate and rotate the peptide during K* optimization. **B & C:** The Inverse Alanine Scanning applied to CALP-DPR5. Complementing the Minimum Flexible Set technique, *Inverse Alanine Scanning* addresses peptide:target clashes by mutating all peptide residues modulo a single amino acid to alanine. In (B) & (C), position P^-1^ (Val) is redesigned, and K* is used to mutate all other peptide residues to alanine, as well as to continuously minimize peptide:target sidechain conformations. **B:** Inverse Alanine Scanning structural prediction with the source amino type, valine. As expected, the clashes present in CALP-DPR5 (A) vanish in the Inverse Alanine Scanning peptide (as indicated by the lack of red and pink Molprobity dots). Furthermore, K* has rotated P^-1^ (Val) to point towards the peptide’s C-term, indicating that conformation is preferable. With this rotation, P^-1^ (Val) remains within CALP’s hydrophobic pocket. **C:** An OSPREY-predicted ensemble of the result of the Inverse Alanine Scan mutating position P^-1^ to methionine. CALP V345 and I295 form favorable van der Waals interactions (green and blue Molprobity dots) with multiple conformations of P^-1^ (Met), which is also reflected in the increase in K* score of P^-1^ (Met) compared to P^-1^ (Val). This result indicates that further K* binding affinity optimization can include methionine at P^-1^, and that V345 and I295 should be allowed to flex continuously in CALP-DPR5 design candidates containing mutations at P^-1^.

##### 2.3.1.2 Design Technique 2: Inverse Alanine Scanning

Inverse Alanine Scanning (IAS) complements the Minimum Flexible Set. Whereas the MFS technique computes a conformation space that resolves all clashes, IAS determines single residue mutations on the peptide that may increase binding to the target protein only when the target protein’s nearby residues are provided sufficient flexibility in the conformation space. We call this Inverse Alanine Scanning because this computational technique mutates all the peptide’s residues *except* the residue of interest to alanine, the opposite of the canonical alanine scanning experiment. Further, IAS not only indicates which residues in the target protein must be flexible to accommodate favorable mutations, but it also indicates which protein residues can safely remain rigid because they do not interact with peptide mutations in their vicinity. When IAS identifies a favorable mutant containing residues that do *not* adopt more than one rotamer, then these residues can safely be omitted from the final conformation space. This pruning technique is valuable because it enables specification of a smaller conformational space for the next step in the pipeline, K*-based Mutational Scanning. See Figure 3 (B & C) for a visualization of this technique.

##### 2.3.1.3 Design Technique 3: K*-based Mutational Scanning

A *Mutational Scan* uses K* to systematically mutate a residue in the DPR scaffold to all 19 other amino acids. We specified a K* design implementing a Mutational Scan for each residue in a DPR peptide and defined the conformation space as the union of the set of flexible residues from the Minimum Flexible Set and the Inverse Alanine Scanning steps. In many of our DPR cases, we observed mutations that notably improved the K* score (an approximation to binding affinity, *K_a_*). We then carried these positive mutations forward into subsequent K* designs that permitted multiple simultaneous peptide mutations in order to optimize peptide:target binding. We then further refined that set by removing sequences whose increase in K* score was driven primarily by peptide destabilization, as indicated by a large decrease in the unbound peptide’s partition function value *q_L_* (Appendix A: the K* algorithm). Finally, we analyzed the OSPREY-predicted structures of the top three sequences for each DPR scaffold.

## 3 Results

### 3.1 DPR validation criteria

To predict novel D-peptides inhibitors, we validated each of the DPR peptides across multiple criteria relevant to PDZ domain inhibitors.

#### The DPR’s binding affinity

An effective inhibitor must interact with its protein target in such a way that it disrupts the target protein’s ability to bind its endogenous ligand. To validate the inhibitory potential of the D-peptides, we compared their K* scores to the K* scores of each PDZ-domain’s endogenous ligand, as well as some previous L-peptide inhibitors. We used the K* algorithm^60,63^ to optimize the D-peptide’s sequence to increase binding affinity between the peptide and the endogenous ligand’s binding site (the groove between α2 and β2 in the PDZ domain).

#### The DPR’s ability to replicate biophysical facets common to PDZ domain binding

Prior research has identified structural and biophysical elements that are commonly found facilitating canonical PDZ domain interactions^11–16,30,83–85^. One such element is the presence of a hydrogen bond network formed between the peptide’s C-terminal carboxylate and the loop connecting β1 and β2, termed the *carboxylate binding loop (CBL)*^12,16,86^. Another is the presence of β-strand-β-strand interactions between the peptide and β2^15,16^. A third commonality is the presence of a hydrophobic pocket in the α2-β2 groove, which canonically is filled by a hydrophobic residue at position P^0^ in the peptide^15^, and in some peptides, by residue P^-2^ ^16^. We assessed the following biophysical facets in our validation of D-peptides: the 1) H-bond network formed by the D-peptide carboxylate and the CBL; 2) β-strand interactions between the D-peptide backbone and β2, and; 3) ability of the D-peptide to fill the hydrophobic pocket. We believe these facets to be sufficient, but not necessary, in the development of novel D-peptide inhibitors (see Section 5 for additional information on this point).

#### The presence of novel and favorable interactions in the DPR

Our validation includes a structural analysis of the OSPREY-predicted low-energy ensemble of the DPR bound to its target PDZ domain. Since DexDesign predicts *de novo* D-peptides, and since empirical structures of D-peptide inhibitors bound to PDZ domains are lacking, it is possible that a D-peptide could bind its target PDZ domain in a mode quite distinct from that of canonical L-peptides. For example, L-peptide residues at positions that point into the PDZ domain’s binding groove may, in a D-peptide, point away (and vice-versa, see Appendix Figure 7 for an example), providing the possibility for some peptide residues to interact with parts of the PDZ domain in ways not formerly possible. To account for the possibility of novel modes of binding (and not, e.g., discard D-peptides that fail to replicate all the criteria above), we analyzed the OSPREY-predicted low-energy molecular ensembles of the DPR bound to its target PDZ domain. In these analyses, we highlight the presence (or absence) of notable structural features capable of further validating the quality of the DPRs.

### 3.2 Designed Inhibitors Targeting CALP

Using our crystal structure of kCAL01 bound to CALP (PDB ID 6ov7)^28^, we used DexDesign to generate 5 DPR scaffolds: CALP-DPR[1-5] (see Appendix Table 1 for further information about the DPR scaffolds). We then applied the design techniques and selection procedures from Section 2.3.1 to optimize the DPRs, thereby obtaining a final set of 15 D-peptide CALP inhibitors, CALP-PEP[1-15]. We assessed each of the CALP-PEPs using the quantitative and structural validation criteria described in Section 3.1. We also compared the CALP-PEPs to CALP’s endogenous ligand (CFTR) and also to the most binding-efficient L-peptide CALP inhibitor, kCAL01, which we reported in 2012^18^ and solved a crystal structure of in 2019^28^. An overview of each of the CALP-PEP’s K* scores, CBL H-bonds, and peptide:β2 backbone H-bonds is shown in Figure 4. Notably, each of the CALP-PEPs is predicted to bind CALP tighter than the CFTR C-terminal SLiM, a critical prerequisite of an effective inhibitor. After normalization (see Appendix B for information on the normalization procedure) and conversion to Gibbs free energy, the top 3 peptides, CALP-PEP9, CALP-PEP4, and CALP-PEP5, when compared to the CFTR C-terminus, have a ΔG of 2.3, 2.3, and 2.1 kcal/mol lower than the CFTR C-terminus (CALP-PEP9: ΔG = -6.9 kcal/mol, CALP-PEP4: ΔG = -6.9 kcal/mol, CALP-PEP5: ΔG = -6.7 kcal/mol, CFTR C-terminus: ΔG = -4.6 kcal/mol), improving K_D_ over the CFTR C-terminus by 46-, 44-, and 33-fold, respectively. Below, we provide the results and analyze the OSPREY-predicted structural ensembles.

**Figure 4:**
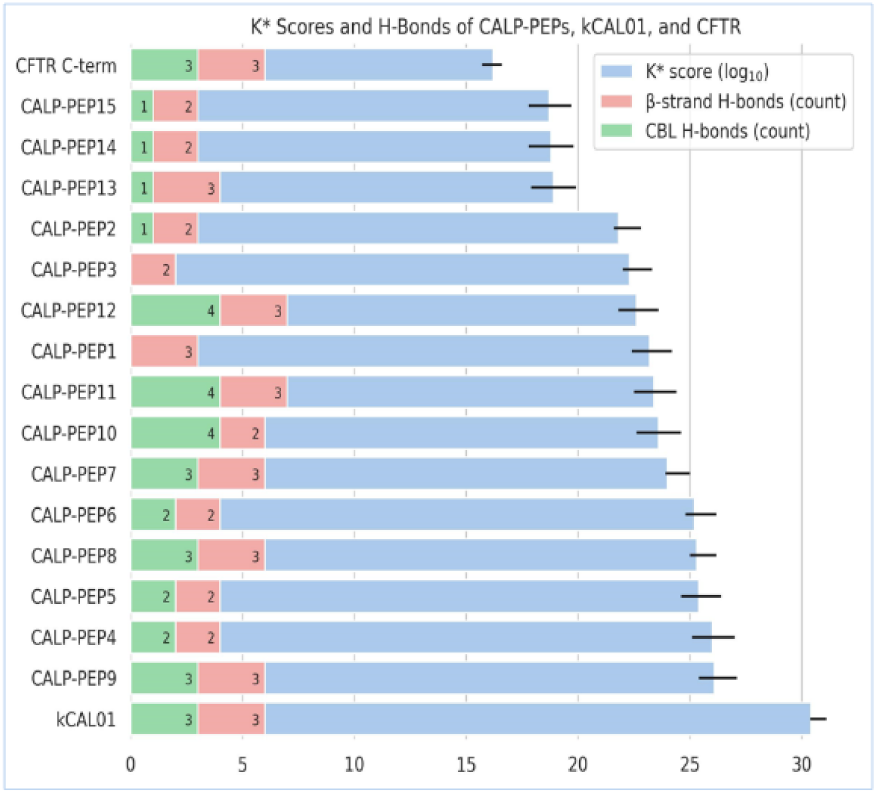
Quantitative and Structural Analysis of the CALP-PEPs, kCAL01, and the CFTR C-terminal SLiM. The OSPREY-predicted binding affinity (log10 K* score) of the tightest known peptide binder of CALP (kCAL01, PDB ID 6ov7)28, the CALP-PEPs, and CALP’s endogenous ligand (the CFTR C-terminal SLiM, PDB ID 2lob)86 show (blue bars) that the CALP-PEPs are predicted to bind more tightly, than the CFTR C-terminal SLiM. Conversely, the CALP-PEPs are predicted to bind CALP less tightly than the best known CALP peptide inhibitor, kCAL0118,28. Since the primary objective of a competitive inhibitor is to outcompete an endogenous ligand in binding to the target protein, the K* scores, *viz.*, provably accurate ε-approximations to *K*a (see Appendix A), of the 15 *de novo* D-peptide CALP-PEPs exceeding that of CFTR’s C-terminal SLiM indicates that the CALP-PEPs meet their fundamental design objective. For example, CALP-PEP9 has a ΔΔG of -2.3 kcal/mol, improving KD 46-fold compared to the CFTR C-terminus (see Appendix B). While not predicted to bind as tightly as kCAL01, D-peptides have therapeutic advantages over L-peptides, including improved metabolic stability, that can compensate for not reaching the binding affinity of the strongest CALP peptide inhibitor. The red bars show the number of β-strand H-bonds contributing to the common β2-sheet extension PDZ-binding motif. The green bars show the number of H-bonds between the peptide’s C-terminal carboxylate and the CBL. The number of CBL and β-strand H-bonds varies across the CALP-PEPs, but CALP-PEP9, the tightest predicted binder, has 3 CBL and 3 β-strand H-bonds, the same number as CFTR and kCAL01. The K* scores of the CALP-PEPs and empirical structures were determined using the K* algorithm41,60 in OSPREY. View Appendix A for a definition of the K* algorithm and K* score. The error bars on the K* scores show the provable upper-and lower-bound of the K* approximation. The number of H-bonds between the peptides and CALP were determined using Pymol62.

K* redesign of the peptide sequence enabled each of the CALP-PEPs to achieve a tighter binding affinity to CALP when compared to the DPR scaffold from which it was generated. This is indicated by their positive log_10_ ΔK* score (Appendix Table 2). Notably, the CALP-PEPs ΔK* scores strongly correlate with their K* scores (Spearman correlation = 0.95). We postulate this strong correlation indicates that DexDesign’s K* optimization is not merely alleviating clashes in the DPR scaffolds, but that it is also identifying peptide sequences forming novel side chain interactions that increase binding affinity. For example, CALP-PEP9’s P^-2^ arginine reaches across β2 and forms a salt bridge with β3’s Glu309 (see Figure 5, D). The magnitude of the predicted improvement in binding ranges from ΔΔG = -2.0 kcal/mol for CALP-PEP15 to -3.5 kcal/mol for CALP-PEP9.

**Figure 5:**
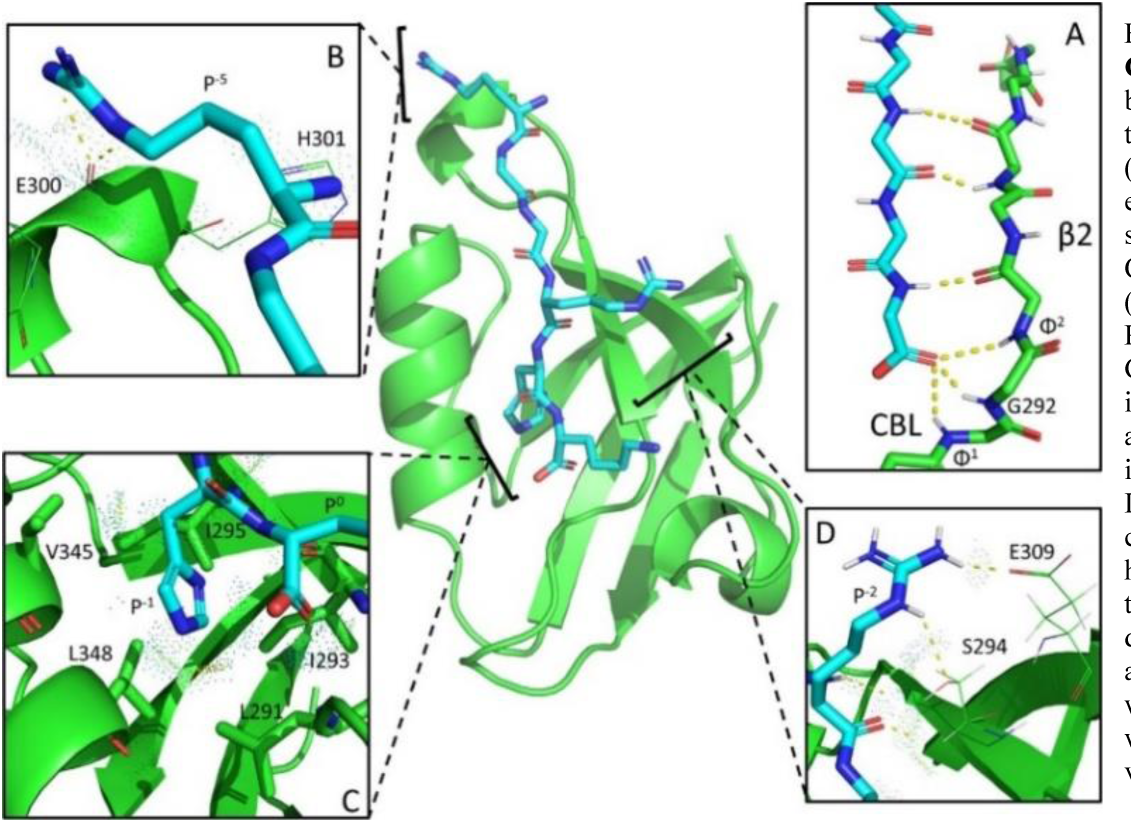
Structural analysis of OSPREY-generated ensemble of CALP-PEP9. Of the 15 CALP-PEPs, CALP-PEP9 (RGGRHK) is predicted to be the tightest binder to CALP with a log10 K* score of 26.1, which approaches the predicted affinity of the most binding-efficient L-peptide inhibitor kCAL01 (previously reported18 by our lab; with a log10 K* score of 30.4), and vastly exceeds the predicted binding affinity of the CFTR C-terminal SLiM (log10 K* score of 16.2). CALP-PEP9 is predicted to improve KD by 46-fold over the CFTR C-term, with a normalized KD of 8.9 μM versus 420 μM87 for the C-terminal SLiM (The normalization procedure is described in the Appendix B). **A**: CALP-PEP9’s P0 carboxylate forms favorable H-bonds with the carboxylate binding loop (CBL: G290-I293, GΦ1GΦ2) and strand β2, mimicking canonical PDZ binding interactions16 of L-peptides. **B**: The amino acid at position P-5 in CALP-PEP9 is arginine. P-5’s amino group and sidechain make favorable van der Waals contacts, indicated by blue and green MolProbity dots63,64,80, with CALP’s E300 and H301. Its guanidino sidechain also forms H-bonds with E300’s carboxyl group. **C**: In canonical L-peptide’s, a PDZ-domain’s hydrophobic pocket is filled by a hydrophobic amino acid at position P0.16 In contrast, all of the CALP DPRs fill the pocket with the amino acid at position P−1. CALP’s hydrophobic pocket, defined as the groove between α2 and β2 and involving V345, I295, I293, L291, and L348, is filled by a histidine in position P-1. **D**: CALP-PEP9’s P-2 is arginine, which is predicted to make favorable van der Waals contacts and form H-bonds with β2’s S294 and the sidechain of β3’s E309. K* scores and additional structural validation of all the CALP-PEPs can be found in Appendix Table 2.

The three canonical PDZ domain binding motifs we used as criteria to further validate the CALP-PEPs were: 1) the presence of an H-bond network between the peptide’s C-terminal carboxylate and the CBL; 2) the presence of peptide:β2 backbone interactions; and 3) whether the D-peptide filled the hydrophobic pocket typically filled by P^0^ in L-peptides. To quantify (1), we counted the number of H-bonds formed between the peptide’s C-terminal carboxylate and the CBL, and to quantify (2) we counted the number of H-bonds between the peptide and β2 backbones. We used visual inspection with MolProbity Probe Dots^79,80^ in our lab’s Protein Design Plugin^81^ to perform a binary classification for (3), more specifically, we classified the hydrophobic pocket as *filled* if a D-peptide’s sidechain contacted the residues within the pocket. As a point of reference, kCAL01 and the CFTR C-terminal SLiM each have 3 H-bonds with the CBL, 3 H-bonds between the peptide and β2 backbone, and fill CALP’s hydrophobic pocket with their P^0^ amino acid (valine for kCAL01, leucine for CFTR C-terminal SLiM). 9 of the 15 CALP-PEPs (CALP-PEP[4-12]) had 2 or more H-bonds between their C-terminal carboxylate with the CBL. CALP-PEPs derived from the CALP-DPR1 and CALP-DPR5 scaffolds had 1 (or in the case of CALP-PEP1 and CALP-PEP3, 0) H-bonds.

Encouragingly, all the CALP-PEPs formed at least 2 backbone H-bonds with CALP’s β2 strand (see Figure 4. Appendix Table 2 includes further H-bond measurements). CALP-PEP9, which we predict to be the tightest binder to CALP, forms 3 C-terminal carboxylate H-bonds with the CBL and 3 backbone H-bonds with CALP’s β2 strand, matching the numbers formed by both the CFTR C-terminal SLiM and kCAL01 (see Figure 5). CALP-PEP9 contains arginines at position P^-2^ and P^−5^, both of which are predicted to make favorable van der Waals contacts with CALP (see Figure 5, B & D). In addition, CALP-PEP’s P^-2^ Arg is predicted to form a salt bridge with E309 and hydrogen bond with S294, and P^-5^ Arg H-bonds with E300 and H301 (Figure 5, B & D). While it does not appear that the quantity of CBL and backbone H-bonds drives the predicted strength of binding of CALP-PEPs (see Figure 4), the CBL does play an important role in determining peptide specificity,^11^ therefore we regard evaluation of D-peptide:PDZ domain H-bonds as necessary components of a larger ensemble of criteria. In contrast, whether a CALP-PEP fills CALP’s hydrophobic pocket is important to designing a tight binder.

All the CALP-PEPs saw an increase in their K* scores when they mutated P^-1^ to an amino acid capable of filling the pocket (see Appendix Table 2). 11 of the 15 CALP-PEPs mutate P^-1^ to histidine, 3 of the 15 to phenylalanine, and CALP-PEP15 is unique with methionine. For example, the mutation to histidine at P^-1^ in CALP-PEP9 fills and favorably interacts with multiple residues in the hydrophobic binding pocket (see Figure 5, C).

### 3.3 Designed Inhibitors Targeting MAST2

Using an NMR structure of PTEN bound to MAST2 (PDB ID 2kyl)^76^, we used DexDesign to generate 3 DPR scaffolds: MAST2-DPR[1-3]. From these 3 DPR scaffolds, we used the design techniques described in Section 2.3.1 to generate 15 peptides, MAST2-PEP[1-15]. Appendix Figure 3 provides an overview of the MAST2-PEPs K* scores and how they compare to MAST2’s endogenous ligand PTEN. Additional structural information about the MAST2-PEPs, such as which residue fills MAST2’s hydrophobic cavity, is available in Appendix Table 3.

PTEN binds MAST2 209-fold tighter than CFTR binds CALP (K_D_ = 1.9 ± 0.05 µM vs. 420 ± 80 µM)^87,88^ and binds MAST2 as tightly as the strongest known inhibitor of CALP, kCAL01 (K_D_ = 2.3 ± 0.2 µM)^18^. In other words, to design competitive inhibitors of the MAST2:PTEN interaction requires us to design D-peptide inhibitors with a stronger affinity than the tightest known L-peptide inhibitor of CALP. Despite the challenge inherent in disrupting the MAST2:PTEN interaction, all the MAST2-PEPs are predicted to bind MAST2 with affinities surpassing PTEN (see Appendix Figure 3). The log_10_ K* scores of the MAST-PEPs range from a low of 29.4 for MAST2-PEP9 to a high of 32.7 for MAST2-PEP4. MAST2-PEP4 is the best DexDesign-generated inhibitor and is predicted to bind MAST2 with a normalized Gibbs free energy ΔG of -8.8 kcal/mol, a -1.1 kcal/mol improvement over MAST2:PTEN, resulting in a 5-fold improvement in K_D_. In some cases a 5-fold improvement might be considered small, but we have previously shown^18,40^ that differences of this magnitude can have profound effects on biological activity. For example, kCAL01 binds only 6-fold tighter than previous competing peptides, such as iCAL35, that were discovered via high-throughput SPOT arrays^18^. However, in *ex vivo* assays (see Section 1.2.1) iCAL35 had non-significant biological activity^18^, whereas the 6-fold tighter binding kCAL01 had significant biological activity. Since the SLiMs modulate a delicate network of competing affinities and specificities^11,12,87^, 5-7x improvements in affinity (such as that achieved by CALP-PEP4) can make the difference between failure and true biological activity.

The MAST2-PEPs replicate some of the canonical L-peptide PDZ-binding motifs, such as the residue in position P^0^ filling the hydrophobic pocket between the PDZ domain’s α2 helix and β2 strand. 9 out of 15 MAST2-PEPs have residue P^0^ filling the hydrophobic pocket (see Appendix Table 3). This contrasts with the CALP-PEPs, where in all cases the residue at position P^-1^ filled the hydrophobic pocket. In the best predicted inhibitor, MAST2-PEP4, the P^0^ leucine fills MAST2’s hydrophobic cavity formed by Tyr17, Phe19, Val77, Ile79, and Leu81 (Appendix Figure 4, C & D). In contrast to PTEN’s P^0^ valine, MAST2-PEP4’s P^0^ leucine forms favorable interactions with all 5 of the cavity’s hydrophobic residues. In addition, a rotation of MAST2-PEP4’s C-terminal carboxylate alleviates a steric clash with the carboxylate binding loop present in the MAST2:PTEN complex.

The MAST2-PEPs also exploit novel geometric features of D-peptides not available to their L-counterparts. For example, MAST2-PEP4’s P^-3^ glutamate makes favorable van der Waals contacts with MAST2’s His73 imidazole side chain (Appendix Figure 4, A). PTEN does not make the analogous interaction, and instead PTEN’s P^-3^ isoleucine is oriented towards MAST2’s β2 strand, and the residue nearest to His73 is a glutamine at P^-4^, whose amide fails to make van der Waals contacts with His73 (Appendix Figure 4, B). The creation of novel favorable interactions with MAST2 is common in the designed MAST2-PEPs and compensates for the loss of some of the canonical PDZ-domain binding motifs.

### 4 Computational Complexity

DexDesign includes several design techniques that compute a K* score from thermodynamic ensembles, which is computationally expensive. Previous sublinear K* algorithms include^64,65^ BBK* with MARK*, but these fail to adequately reduce the number of partition function calls for D-peptide redesigns enumerating a large search space. To reduce the number of calls to K* by pruning prohibitively large sequence spaces, we implemented and analyzed the following techniques.

#### DPR Scaffold Generation

Starting with a given L-peptide:L-target complex, steps 1-3 (see Figure 2) of the DexDesign algorithm output a D-peptide query (*Q*) for input to the MASTER search algorithm. While the MASTER algorithm is worse-case exponential in the number of disjoint query segments^59^, in DexDesign *Q* is a single segment and therefore the time required to calculate the optimal rotation and translation matrix with the Kabsch algorithm^89,90^ between *Q* and each contiguous, equally sized segment in a MASTER database with *s* residues is O(*s*). MASTER can compute over a million Kabsch superimpositions per second^59^, leading to empirical runtimes on the order of seconds. MASTER returns the *u* best results in order of backbone RMSD, so DexDesign generates *u* DPR scaffolds, with each scaffold containing a D-peptide *P* and a protein target *T*.

#### Minimum Flexible Set (MFS)

In contrast to the upper bound time complexities given for other redesign methods, MFS provides a lower bound on the size of the conformation space that must be searched or pruned by the downstream K* design. The MFS lower bound predicts the running time of the K* designs and can be used to eliminate infeasible designs. In this sense, MFS serves a role similar to TESS in BWM* as an efficiently computable metric of problem complexity that predicts designability^91^ (see Jou et al., 2016 for TESS proof).

Assume *P* has *n* residues and *T* has *r* residues. By computing the distances between the atoms in *P* and *T*, the Minimum Flexible Set can be computed in O(*nr*) time. *P* is often much smaller than *T* and, in such cases, it is more efficient to compute a bounding ball around the peptide inflated by 5 Å in O(*n*) time and clip *T*’s atoms to those in the ball in O(*r*) time. Let *d* be the number of *T*’s residues in the ball, then the Minimum Flexible Set can be computed in O(*r* + *nd*) time. The Minimum Flexible Set is comprised of *c* clashing peptide residues. We prune DPR scaffolds where the ratio of *c* to *n* exceeds 3/4, reducing the number of DPR scaffolds *u* for which we compute Inverse Alanine Scanning and K*-based Mutational Scans. Furthermore, *c* contributes to the lower bound on the size of the conformation space required to be searched or pruned by the K*-based Mutational Scans. Let *r* be the minimum number of rotamers for a clashing residue, then the size of the conformation space input to the K* search is O(*r*^*c*^). This is important because in practice the time required to compute an ε-accurate partition function for a protein sequence is dependent on the size of the conformation space^64,65,92^, so Minimum Flexible Set aids in pruning DPR scaffolds for which partition functions would be challenging to compute. Empirically, *c* averages 2.0 for kCAL01 and 3.7 for MAST2.

#### Inverse Alanine Scanning & K*-based Mutational Scan

For each of the *n* residue in *P*, Inverse Alanine Scanning mutates the amino acid at that residue to the 19 other amino acids, while mutating all other peptide amino acids to alanine. This generates a total of 20*n* sequences, for which DexDesign computes K* scores. Inverse Alanine Scanning limits the size of the conformation space, *k*, by limiting the flexible residues to *P*’s mutating residue and nearby residues on *T*; in practice the median *k* for CALP was 3,705 conformations and for MAST2 8,580 conformations. A mutation that Inverse Alanine Scanning predicts to ablate peptide:target binding affinity is pruned from further consideration in the K*-based mutational scan. With an amino acid library of size *a*, the number of possible peptide sequences is *a*^*n*^. In contrast, Inverse Alanine Scanning can reduce the sequence space to (*a* − *R*)^*n*−*r*^, leading to an exponential reduction in the number of sequences. In the case of CALP and MAST2, *R* and *r* are on average 16.2 and 2.6, respectively (see Appendix Figure 9 for empirical runtimes), and DexDesign reduces the number of possible peptide sequences by a factor of 1.4 x 10^-4^. By using bounded partition function3 sizes and sparse residue interaction graphs^61,91^, we can compute the K* score in time 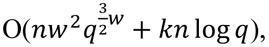 where *w* is branch width, *q* is the number of rotamers per residue, and *k* is number of conformations in a partition function. When *w* is O(*1*), this is polynomial time^91,93^. For the DexDesign experiments we describe, we found that our ε-accurate algorithms^51,94,95^ completed with a median time of 5.4 minutes over all designs and were therefore fast enough for a pre-clinical pipeline.

## 5 Discussion and Conclusions

Although the CALP-PEPs and the MAST2-PEPs both target PDZ domains, the interactions they make with their respective targets can be generally categorized along two quantitative axes: replication and restitution*. Replication* measure reproduction or imitation of interactions previously observed in L-peptide inhibitors, for example, in the case of the PDZ domains, the β2 strand extension, residue P^0^ filling the hydrophobic pocket, and the H-bond network a peptide’s terminal carboxylate makes with the CBL. *Restitution*, on the other hand, measures the process of compensating for typical L-peptide binding motifs by forming novel interactions that are now possible to explore due to the change from L-to D-chirality of the peptide. Whereas in some cases we achieve improved binding through replicating the canonical PDZ interactions, in other cases, due to the special geometric properties of D-peptides, we instead observe an increase in binding (restitution) due to novel interactions that we observe in the structures that were not available to L-peptides. This suggest the intriguing possibility that some peptides may be stabilized by replicating native-like interactions from L-peptides, whereas others might be stabilized by forming novel interactions, available only to the ligands with D side chains. In the *de novo* peptides we generated, both the CALP-PEPs and the MAST2-PEPs contained elements both replicating and restituting the binding interactions formed by the endogenous L-peptides from which their backbone conformations are derived (see Figure 5). Appendix C includes a thorough discussion on replication and restitution for CALP and MAST2. Experimental wet lab data is desirable to increase the credibility of these redesigns and is a valuable future endeavor. Nevertheless, here, to increase confidence in the computed K* scores, we have included a direct comparison between computed K* scores and experimentally measured binding affinity for three peptides (Appendix Table 7).

We also performed an experiment to measure the ability of DexDesign to design a *de novo* D-peptide similar to a D-peptide found in an empirical D-peptide:L-protein complex. With this validation experiment, we report that DexDesign generates a D-ligand with chemistry unique to the DPR scaffold, as opposed to replication of interactions. Further, we also performed a control experiment to investigate the predicted binding of the empirical D-peptide sequence on the MASTER-generated backbone. Interestingly, a 100% wildtype sequence recovery mutant yields lower predicted binding despite the selection of a DPR scaffold with the lowest backbone RMSD. Overall, this experiment highlights the utility of DexDesign for generation of novel peptides, as opposed to sequence recovery of known binders. See Appendix Section D for a full report on this experiment.

In this paper, we presented a new algorithm, DexDesign, to computationally design *de novo* D-peptides. We also presented three new protein design techniques, the Minimum Flexible Set, Inverse Alanine Scan, and K*-based Mutational Scan, that are generally applicable to both D-and L-peptide design. DexDesign required us to add new capabilities to the OSPREY^35^ protein redesign software, including the ability to add arbitrary conformation libraries. Future designs and algorithms that model non-proteinogenic molecular building blocks, such as non-canonical amino acids or small molecule rotamers, are now substantially easier to realize. A comparison study of other computational design programs to OSPREY to further contextualize DexDesign in the redesign community is desirable and is a valuable future endeavor (see Appendix E for more information).

We used DexDesign to generate and optimize 30 *de novo* D-peptide inhibitors for two biomedically important PDZ targets: CALP and MAST2. We computed provable approximations of binding affinity and analyzed the OSPREY-predicted low-energy ensembles of the bound D-peptide:target structures to assess the quality of the novel peptides. We employed a novel restitution-replication framework for analyzing the basis upon which our DexDesign-generated D-peptides improved binding compared to their targets’ endogenous ligands. DexDesign is not restricted to PDZ-domains, it can be applied to design novel antineoplastic, antifungal, antiviral, or antimicrobial D-peptide therapeutics. It is a general algorithm applicable to any target for which there exists structural models of a peptide:target complex. The structural models can be determined experimentally or computationally predicted using machine learning-based algorithms such as AlphaFold^96,97^, although the accuracy of the results may be somewhat diminished compared to experimentally determined structures of ligand-target complexes. Thus, DexDesign provides an important tool to the drug discovery community for developing novel D-peptide therapeutics.

## Appendix

To substantiate the claims of the main paper, and to provide full details of our methods and results, this appendix provides additional information and data. Section A begins with a review of the K* algorithm, which is implemented in DexDesign as introduced in Section 1.4 of the main paper. Section B includes a discussion on normalization of K* scores and conversion to free energy. Section C includes examples of replication and restitution for MAST2 and CALP. Section D describes a validation experiment for DexDesign on the peptide GyGlanvdessG. Section E includes an analysis of validation methods for computational tools. Section F displays supplementary tables on the DPR scaffolds, K* scores, structural features validation for CALP and MAST2, the GyGlanvdessG validation experiment, and Spearman rank correlation coefficient experiments. Finally, we conclude the appendix with figures in Section G, which includes, in order, the reflection operation for peptides, OSPREY GUI screenshots, MAST2 DPR results, MAST2-PEP4 restitution, properties of DPRV (validation experiment), the composition of the MASTER database, orientation differences between L and D peptides, backbone orientation comparisons in the GyGlanvdessG validation experiment, DexDesign runtimes for CALP and MAST2, and Spearman rank correlation data for cRAF-RBD.

### A. The K* algorithm

Here, for the reader, we review the K* algorithm in the OSPREY protein design software suite^1^, which we have presented and analyzed in Lilien et al. (2005)^2^, Georgiev et al. (2008)^3^, Donald (2011)^4^, Gainza et al. (2012)^5^, Hallen et al. (2018)^1^, Ojewole et al. (2018)^6^, and Jou et al. (2020)^7^. In brief, the K* algorithm computes a provably good ε-approximation to the binding affinity constant, *K*_a_. K* does so by calculating an ε-accurate partition function for three structures: the bound protein:ligand complex (denoted *PL*), the unbound protein (denoted *P*), and the unbound ligand (denoted *L*). Let *X* be an arbitrary state, *X* ∈ { *P*, *L*, *PL* }. The partition function is the sum of the Boltzmann-weighted energies for all the conformations in the thermodynamic ensemble of *X*. This encodes the physical fact that low-energy conformations are more likely than high energy conformations, and are therefore weighted more. Let *S* denote an arbitrary amino acid sequence, then the partition function of *S* in state *X* (which we denote as *q_x_(s))* is:

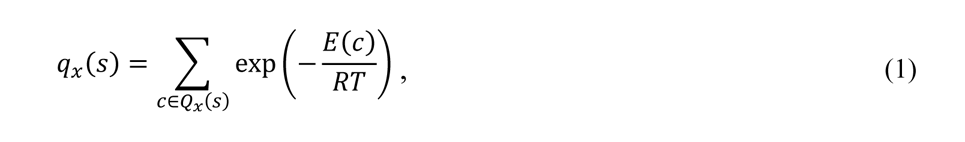

where *Q*_*x*_(*S*) is the entire conformational ensemble of sequence *S* in state *X*, and *c* is a single conformation from that ensemble. *E*(*c*) is the energy of conformation *c*. *R* is the ideal gas constant and *T* is the temperature in absolute Kelvin.

The K* score for a sequence *S* approximates *K*_*a*_:

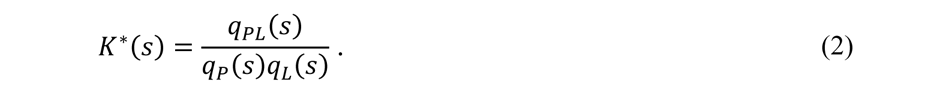

K* uses minimized continuous rotamers when computing *E*(*c*), as described by the iMinDEE algorithm^5^. It uses the A* algorithm^8,9^ to search over *Q*_*X*_(*S*) and streams a gap-free list of conformations in order of the lower-bound of a conformation’s minimized energy^5^. In this fashion, while computing a running sum that adds up the non-decreasing sequence of statistical weights, every future conformational energy will be higher, and therefore will have a smaller weight. By this reasoning, and as reported previously ^10–15^, the tail of the Boltzmann sum is bounded which enables us to obtain a good epsilon approximation algorithm for the partition function in the computational model. It then minimizes the conformation to calculate *E*(*c*), and generates an ε-approximation of the partition function *q*_*X*_(*S*) and the ensemble-complete K* value. This approximation is known as the K* score.

### B. K* Score Normalization

The K* algorithm^2,5^ and the K* scores it predicts have been validated experimentally in many previous works^5,16–23^. While sometimes we have observed the K* scores to correlate quantitively (Pearson) with^15,18,20^ K_a_, we have greater evidence that K* scores better correlate with K_a_ using a ranking (Spearman) paradigm^1,23^. For example, we recently analyzed the accuracy of EWAK* (an accelerated K*-derivative algorithm) predictions on 41 c-Raf-RBD variants binding to KRas and found that the K* score and experimental ranks correlated with a Spearman ρ of 0.81^23^. This data is provided in Appendix Table 5 and visualized in Appendix Figure 10. We also evaluated OSPREY predictions across four protein systems^15,24^, producing an “across all” Spearman rank correlation coefficient of 0.762. This was calculated by ranking each system individually and then calculating Spearman’s across all designs (Appendix Table 6). Therefore, the Spearman rank correlation coefficient is sufficient for rank-based protein screening and provides accurate binding affinity guidance for experimental labs testing candidates in order of predicted ranking of binding affinity. Therefore, when we convert K* scores to physical quantities, such as ΔG, we normalize the K* scores based on available empirical evidence (when available). For example, for the CALP-PEPs, we scale our ΔG values based on existing binding affinity data for CFTR (K_i_ = 420 ± 80 μM)^25^ and kCAL01 (K_i_ = 2.3 ± 0.2 μM)^21^, for which we have also predicted K* scores (see Appendix Table 2).

The necessity to normalize K* scores when converted to physical quantities is due to simplifications necessary for modeling the systems computationally. We have previously described normalization in Wang (2022)^26^, and review the reasons here for the reader. First, since it is computationally intractable to model large continuous movements of each atom in a molecule while simultaneously searching over protein sequences and conformations, our K* computations focus on modeling continuous movements for residues within the PDZ binding site. Second, the OSPREY energy function models solvent using a residue-pairwise approximation to the energy field, namely the Effective Energy Function (EEF1) for proteins in solution^27^. Analysis of previous K* designs has shown that the EEF1 contribution to the OSPREY energy function can overestimate van der Waals terms. Lastly, limitations in the input model, as well as the user-specified conformation space can cause an over-or underestimation in the K*-predicted enthalpy or entropy of the bound protein:peptide complex.

### C. Replication and Restitution in CALP and MAST2

Consider the SLiM’s canonical C-terminal carboxylate H-bond network formed with the CBL. Whereas 13 of the 15 CALP-PEPs replicated (to varying degrees, and sometime even exceeding the number of) H-bonds formed with the CBL, the MAST2-PEPs’ terminal carboxylate tended to have few, if any, H-bonds with the CBL (see, e.g., Appendix Figure 4, C). We postulate that the reason for the MAST-PEPs’ use of restitution instead of replication for the CBL H-bond network is that the structure of the PTEN:MAST2 complex (PDB ID 2kyl)^28^ indicates the existence of steric clashes between the C-terminal carboxylate and the CBL (see Appendix Figure 4, D). When using MolProbity^29^ to evaluate the lowest-energy model of the empirical NMR structure, it indicates there is a bad clash (van der Waal radii overlap of 0.518 Å) between PTEN P^0^ valine’s OXT atom and the HA atom from Lys16 in MAST2’s CBL. OSPREY’s energy function^1,5,30,31^, which uses continuous sidechain minimization in addition to translation and rotation of the peptide to minimize the energy of each conformation evaluated by OSPREY’s iMinDEE/K* algorithm^3,5^, pushes and rotates the C-terminal carboxylate away from the CBL to alleviate the steric clash (see Appendix Figure 4, C), with the trade-off being the loss of replication of some of canonical PDZ CBL H-bond interactions.

The CALP-PEPs and MAST2-PEPs also exploit restitution to form novel favorable interactions with their target proteins. One type of restitution is the creation of favorable new side chain interactions between the peptides and their targets. For example, CALP-PEP9’s P^-2^ arginine forms one H-bond and a salt bridge with CALP β2 strand’s S294 and the sidechain of β3’s E309 (see Figure 5, D) that are absent in the CALP:kCAL01 structure (PDB ID 6ov7)^22^. Interestingly, CFTR’s P^-1^ arginine forms an analogous H-bond with E59 CALP:CFTR structure (PDB ID 2lob)^32^, providing evidence that this interaction, as restituted in CALP-PEP9 (based on the kCAL01 structure lacking this interaction), is both plausible and favorable *in vitro*. The MAST2-PEPs likewise restitute novel favorable interactions. For example, MAST2-PEP4’s P^-3^ glutamate makes favorable van der Waals contacts with MAST2’s His73 imidazole side chain (see Appendix Figure 4, A). In contrast, PTEN cannot make some favorable interactions available to MAST2-PEP4. In the MAST2:PTEN structure, PTEN’s P^-3^ isoleucine is oriented towards MAST2’s β2 strand, and the residue nearest to His73 is P^-4^ is glutamine, whose amide fails to make van der Waals contacts with His73 (see Appendix Figure 4, B). In the future, we believe that designed D-peptide libraries of binders and inhibitors can be characterized as falling on a spectrum ranging from replication (1) to restitution (-1) and can be visualized as a per-residue replication-restitution score ranging from 1 to -1. In this way, the functional contributions to binding of *de novo* peptides could be mapped into a vector space which can be visualized or exploited as novel features for machine learning design approaches.

### D. Validation of DexDesign scaffold discovery and redesign

To assess DexDesign, we performed an experiment to measure the ability of DexDesign to design a *de novo* D-peptide similar to a D-peptide found in an empirical D-peptide:L-protein complex. In our experiment, we began with the crystal structure of a known D-peptide in complex with an L-protein and applied a global reflection to the D-peptide:L-protein complex. Then, we employed MASTER to search the L-database using the resulting reflected L-peptide as the query. The returned L-structures were aligned with the reflected complex and reflected once again to produce D-peptide:L-protein scaffolds ordered by increasing backbone alignment RMSD. The first, and therefore lowest, RMSD backbone alignment was then selected for redesign using OSPREY. We then measured the similarities of the redesigned D-peptide to the empirical D-peptide. While DexDesign is intended for construction of novel *de-novo* D-peptides, the capability to generate a DPR similar to the D-form empirical structure should validate our redesign protocol.

We selected a crystal structure of the D-amino acid containing peptide GyGlanvdessG in complex with streptavidin (PDB ID 5n8j)^33^. Streptavidin is a homotetrameric protein that binds the vitamin biotin with high affinity^34^, and is therefore commonly used in Western blotting and immunoassays^35^. A monomer of streptavidin forms a β barrel, with ligands oriented towards the interior of the barrel. Similar to CBL interactions with CALP and MAST2, streptavidin forms favorable hydrogens bonds with ligands via a flexible binding loop^36^. Analogously, streptavidin exhibits hydrophobic contributions through inward-facing tryptophan residues of the β barrel^37^. Therefore, a high-affinity ligand should establish hydrogen bonds with the binding loop while orienting hydrophobic residues towards β barrel tryptophans. We selected this system due to its comparable D-peptide size and similar chemistry to PDZ domains.

We sourced the lowest backbone RMSD (0.48 Å) of inverted D-amino acid GyGlanvdessG from chain A residues 608 to 616 of ST0929 (PDB ID 3hje)^38^, a glycol transferase. After application of Minimum Flexible Set, Inverse Alanine Scanning, and K*-based Mutational Scan techniques to this scaffold, we determined the optimal binder, hereafter denoted as DPRV, to have a log_10_ K* score of 32.8 with the sequence WWMIGDWND. This differed slightly from GyGlanvdessG (GLANVDESS), which has a log_10_ K* score of 32.2. The sequence similarity between the two peptides is 21.43%, a degree of native sequence recovery comparable to reported recovery in popular protein design programs such as Rosetta for L-proteins^39^. This is especially true for NMR structures, such as we used in our MAST2 study (see Section 3.3). With DPRV, we report that DexDesign generates a D-ligand with chemistry unique to the DPR scaffold.

While DexDesign exhibits comparable performance to state-of-the-art methods^39^, native sequence recovery on a short (9 residue) peptide may be a poor indicator of ligand binding. For example, a 40% sequence recovery equates to 3.6 residues for our redesigned peptide (for further discussion on native sequence recovery, see Appendix E). This is a small number of residues, and likely fails to capture the geometric and chemical features that drive high affinity. To investigate the similarities of GyGlanvdessG and DPRV, we also report the backbone alignment RMSD of DPRV to GyGlanvdessG:streptavidin (0.48 Å), and the geometric, chemical, and biophysical properties of our designed peptide that enable binding (see Appendix Figure 5). Finally, we report the log_10_ K* scores computed over molecular ensembles as validation of binding competency (above and see Appendix Table 4).

We also performed a control experiment wherein we mutated the ST0929 scaffold sequence (RYEEGLFNN) directly to the sequence of the D-peptide GyGlanvdessG, without using any OSPREY-based techniques such as K*-based Mutational Scan and Inverse Alanine Scanning. The purpose of this experiment was to investigate the predicted binding of the GyGlanvdessG sequence on the ST0929 scaffold backbone. This control experiment produced a log_10_ K* score of only 26.6, a difference of -5.7 from GyGlanvdessG. Interestingly, a 100% wildtype sequence recovery mutant yields lower predicted binding despite the selection of a DPR scaffold with the lowest backbone RMSD. Therefore, we conclude that the GyGlanvdessG sequence was not recovered by the full DexDesign protocol because these novel OSPREY-based design techniques would not permit optimal binding on the lowest backbone RMSD scaffold. Instead, the DexDesign techniques (as outlined in Section 2.3.1) resulted in a novel D-peptide. These results highlight the sensitivity of the peptide design to the starting scaffold; even similar-appearing scaffolds have different degrees of designability due to geometric differences between backbones. Overall, our experiment highlights the utility of DexDesign for generation of novel peptides, as opposed to sequence recovery of known binders.

The difference between amino acid composition of the D-amino acid containing peptide GyGlanvdessG and the redesigned peptide is likely due to subtle differences in scaffold geometry. As shown in Appendix Figure 5 and 8, the backbones of D-amino acid GyGlanvdessG, DPRV, and the ST0929-sourced peptide scaffold mutated to the endogenous ligand (control) vary at residues important for establishing hydrogen bonds. For example, GyGlanvdessG’s Glu7 residue makes hydrogen bonds with residues Asn23 and Ser27 of streptavidin (Appendix Figure 8, A). DPRV’s Trp7, which is shifted 1.9 Å away from streptavidin Ser27 in comparison to GyGlanvdessG, does not form either of these hydrogen bonds. However, DRPV’s Asp9 facilitates hydrogen bond formation with residues Ser45 and Ser52 on streptavidin (Appendix Figure 5, B). These residues belong to the flexible binding loop, where favorable contacts are crucial for high-affinity binding^34^. The hydrogen bond formed with flexible loop residue Ser52 is unique to DRPV and is not present in the ST0929 control experiment or GyGlanvdessG. Therefore, a peptide that replicates some characteristics of a known ligand, while restituting novel interactions, may be a much more competent binder.

### E. Comparison of Methods and Native Sequence Recovery

While empirical comparison of structural prediction algorithms is sound, empirical comparison of the output of design algorithms is more complicated due to conformational epistasis. Namely, many sequences can fold to the same backbone conformation. This is necessary for evolution because it’s important that spontaneous mutations do not always disrupt the fold; this allows the evolution of new function. This is why high native sequence recovery is undesirable: a 25% to 50% native sequence recovery is considered excellent, and higher is considered overfitting^40^. In the case of native sequence recovery, the community has converged on this desired percentage^41^, which is empirically grounded, because computational predictions are being compared to experimental sequences. However, issues arise when making direct comparisons between computational predictions.

The issue that arises is that protein design algorithms, in general, are not *complete*. They may return some sequences with the desired structure and function, but they are not guaranteed to return all such possible sequences. Thus, in principle, two sound algorithms can return two different sets of sequences and those sets may, or may not, overlap. If a sequence is missing from one set, it does not necessarily mean the algorithm is wrong. The primary validation would be experimental analysis of folding, structure, and binding. For this reason, an end-to-end, purely computational comparison of *in silico* design methods is methodologically fraught.

An example is our paper, in *Cell* 2023^42^, which reports the computational design of three antibodies against HIV. The antibody similar to PG9 that we report, called DU011 (PG9-RSH N(100f)Y), contains four mutations, one of which had previously been discovered by Rosetta^43^. This could be viewed as a form of “validation” in that OPREY returned a similar mutant to that predicted by Rosetta. On the other hand, Rosetta also predicted sequences that did not work. OSPREY did not predict these. Finally, OSPREY predicted sequences that Rosetta did not, in particular the Duke novel antibody DU025, which is broader and more potent than DU011. In the paper, we measured the neutralization breath and potency and solved the cryo-EM structures to demonstrate this binding mode and mechanism of action. Therefore, solely comparing the computational predictions from Rosetta and OSPREY would likely result in some overlap, but also sets that are largely disjoint. Without experiments, we would not be able to definitively say which is correct, and certainly in this case, we couldn’t say that Rosetta was ground truth. This phenomenon was discussed extensively in our 2012 paper^13^.

Finally, even in the discrete rotamer design case, computing the GMEC or performing positive multistate design is NP-Hard^44,45^. This corresponds to finding one solution in an optimization problem. But, if we expect the algorithm to be complete, namely, to return all sequences that have the correct structure and function, this complexity increases to either co-NP-hard or #P-hard. Assuming the usual structural complexity conjectures, this would move the problems into a harder complexity class, and few current protein design algorithms claim to be complete in this sense^45,46^. Nevertheless, extensions of OSPREY, such as the BBK* or MARK* algorithms^10,47^, do guarantee to enumerate conformations and sequences gap-free in order of free energy. Theoretically, this enumeration should be guaranteed to produce all sequences that, within the model, have the desired binding and folding characteristics. However, even if we were to compare the output of a complete algorithm such as OSPREY/BBK*^10,24^ to Rosetta^48^, we are relegated to comparing provably-complete enumeration to a sparse Monte Carlo sampling that is known to be incomplete^49^. This was discussed extensively in our previous papers^13,50^. OSPREY and Rosetta both make predictions that are theoretical, and for the reasons provided, comparing the list of predictions directly presents methodological hurdles for comparison testing.

### F. Appendix Tables

**Appendix Table 1:** The DPR scaffolds for each of the D-peptide redesigns. Each of the DPR scaffolds was extracted from an extant empirical protein structure from the Protein Databank^51,52^. **Name:** the name of the DPR scaffold. **Source PDB ID:** the PDB ID that can be used to retrieve the empirical structure from the Protein Databank. **Source Residues:** The chain and amino acid range in the empirical structure uniquely identifying the source of the DPR. **Template Peptide:** The PDZ-binding peptide used as a search query to MASTER^53^, which returned a result set containing the DPR scaffold. **Backbone RMSD (Å):** the full backbone RMSD between the template peptide and the DPR scaffold. **Sequence:** The DPR scaffold’s peptide sequence.

| Name | Source PDB ID | Source Residues | Template Peptide | Backbone RMSD (Å) | Sequence |
| --- | --- | --- | --- | --- | --- |
| CALP-DPR1 | 1g1k | A117-122 | kCAL01<sup>22</sup> | 0.91779 | DGGAFG |
| CALP-DPR2 | 3u0o | A137-142 | kCAL01 | 0.88123 | AGGHSI |
| CALP-DPR3 | 4m6r | A45-51 | kCAL01 | 0.88412 | GGGISL |
| CALP-DPR4 | 7bjt | B403-409 | kCAL01 | 0.89913 | QGGVAI |
| CALP-DPR5 | 1c8n | C61-66 | kCAL01 | 0.98744 | AGGFVT |
| MAST2-DPR1 | 3s4k | A76-81 | PTEN<sup>28</sup> | 0.91439 | EGGSVV |
| MAST2-DPR2 | 6zec | H5-10 | PTEN | 0.9417 | QQWGAG |
| MAST2-DPR3 | 6tc2 | L22-27 | PTEN | 0.95832 | RGFFYT |

**Appendix Table 2:** K* scores and structural features validation of the CALP-PEPs, kCAL01, and the CFTR C-terminal SLiM. **ID:** the ID of the DexDesign-generated D-peptide inhibitor, or the endogenous ligands. **DPR ID:** The scaffold from which the CALP-PEP was generated (see Appendix Table 1 for the structural source of the DPR) or, in the case of kCAL01 and the CFTR C-term SLiM, the PDB ID of the structure. **Sequence:** the amino acid sequence of the peptide. **K* score (log_10_) [bounds]:** The OSPREY-computed K* score (see Appendix A for a description of the K* algorithm^2,3^) and the computed lower-and upper-bounds. The K* scores were computed to an ε of 0.9. **ΔK* score (log_10_):** The increase in the CALP-PEP’s K* score over its source DPR scaffold, indicating the predicted improvement in binding K* and DexDesign were able to achieve through sequence optimization. **CBL H-bonds:** The number of H-bonds formed between CALP’s carboxylate binding loop (CBL) and the peptide’s C-terminal carboxylate. **β-strand H-bonds:** the number of H-bonds formed between CALP’s β2 strand and the peptide mainchain. **Residue at P^-1^:** The type of amino acid filling CALP’s hydrophobic pocket at the peptide’s position P^-1^ († position P^0^ in L-peptides).

| ID | DPR ID | Sequence | K* score (log<sub>10</sub>) [bounds] | ΔK* score (log<sub>10</sub>) | CBL H-bonds | β-strand H-bonds | Residue at P<sup>-1</sup> |
| --- | --- | --- | --- | --- | --- | --- | --- |
| **KCAL01** | 6ov7<sup>22</sup> | WQVTRV | 30.4 [30.4 - 31.1] | N/A | 3 | 3 | Ile<sup>†</sup> |
| **CFTR C-TERM** | 2lob<sup>32</sup> | VQDTRL | 16.2 [15.7 - 16.6] | N/A | 3 | 3 | Leu<sup>†</sup> |
| **CALP-PEP1** | CALP-DPR1 | RMGRFK | 23.2 [22.4 - 24.2] | 11.1 | 0 | 3 | Phe |
| **CALP-PEP2** | CALP-DPR1 | DMGRFK | 21.8 [21.6 - 22.8] | 9.7 | 1 | 2 | Phe |
| **CALP-PEP3** | CALP-DPR1 | RGGRFK | 22.3 [22.0 - 23.3] | 10.2 | 0 | 2 | Phe |
| **CALP-PEP4** | CALP-DPR2 | RQGRHI | 26.0 [25.1 - 27.0] | 13.0 | 2 | 2 | His |
| **CALP-PEP5** | CALP-DPR2 | AQGRHM | 25.4 [24.6 - 26.4] | 12.4 | 2 | 2 | His |
| **CALP-PEP6** | CALP-DPR2 | AQGRHI | 25.2 [24.8 - 26.2] | 12.2 | 2 | 2 | His |
| **CALP-PEP7** | CALP-DPR3 | RGGIHK | 24.0 [23.9 - 25.0] | 11.2 | 3 | 3 | His |
| **CALP-PEP8** | CALP-DPR3 | RGGRHL | 25.3 [25.0 - 26.2] | 12.5 | 3 | 3 | His |
| **CALP-PEP9** | CALP-DPR3 | RGGRHK | 26.1 [25.4 - 27.1] | 13.3 | 3 | 3 | His |
| **CALP-PEP10** | CALP-DPR4 | RQGRHM | 23.6 [22.6 - 24.6] | 10.1 | 4 | 2 | His |
| **CALP-PEP11** | CALP-DPR4 | QQGRHM | 23.4 [22.5 - 24.4] | 9.9 | 4 | 3 | His |
| **CALP-PEP12** | CALP-DPR4 | RQGVHM | 22.6 [21.8 - 23.6] | 9.1 | 4 | 3 | His |
| **CALP-PEP13** | CALP-DPR5 | AGGRHR | 18.9 [17.9 - 19.9] | 7.6 | 1 | 3 | His |
| **CALP-PEP14** | CALP-DPR5 | AGGRHM | 18.8 [17.8 - 19.8] | 7.5 | 1 | 2 | His |
| **CALP-PEP15** | CALP-DPR5 | AGGRMK | 18.7 [17.8 - 19.7] | 7.4 | 1 | 2 | Met |

**Appendix Table 3:** K* scores and structural features validation of the MAST2-PEPs and PTEN6. **ID:** the ID of the DexDesign-generated D-peptide inhibitor, or the endogenous ligands. **DPR ID:** The scaffold from which the MAST-PEP was generated (see Appendix Table 1 for the structural source of the DPR) or, in the case of PTEN6, the PDB ID of the structure. **Sequence:** the amino acid sequence of the peptide. **K* score (log_10_) [bounds]:** The OSPREY-computed K* score (see Appendix A for a description of the K* algorithm^2,3^) and the computed lower-and upper-bounds. The K* scores were computed to an ε of 0.9. **ΔK* score (log_10_):** The increase in the MAST2-PEP’s K* score over its source DPR scaffold, indicating the predicted improvement in binding K* and DexDesign were able to achieve through sequence optimization. **Hydrophobic Pocket (location):** The type and location of the amino acid filling MAST2’s hydrophobic pocket. - indicates this residue is absent.

| ID | DPR ID | Sequence | K* score (log<sub>10</sub>) [bounds] | ΔK* score (log<sub>10</sub>) | Hydrophobic Pocket (location) |
| --- | --- | --- | --- | --- | --- |
| PTEN6 | 2kyl<sup>28</sup> | TQITKV | 28.8 [28.4 - 29.8] | N/A | Val (P<sup>0</sup>) |
| MAST2-PEP1 | MAST2-DPR1 | EMGDMD | 30.9 [30.8 - 31.8] | 20.2 | Met (P<sup>-1</sup>) |
| MAST2-PEP2 | MAST2-DPR1 | EEGDMD | 30.5 [30.4 - 31.1] | 19.9 | Met (P<sup>-1</sup>) |
| MAST2-PEP3 | MAST2-DPR1 | EQGDMD | 30.3 [30.2 - 31.0] | 19.7 | Met (P<sup>-1</sup>) |
| MAST2-PEP4 | MAST2-DPR2 | EGEEDL | 32.7 [32.7 - 33.0] | 22.0 | Leu (P<sup>0</sup>) |
| MAST2-PEP5 | MAST2-DPR2 | YGEEDL | 32.6 [32.6 - 32.9] | 21.9 | Leu (P<sup>0</sup>) |
| MAST2-PEP6 | MAST2-DPR2 | FGEEDL | 32.2 [32.2 - 32.3] | 21.5 | Leu (P<sup>0</sup>) |
| MAST2-PEP7 | MAST2-DPR3 | EDDEYE | 30.7 [30.7 - 30.7] | 21.9 | - |
| MAST2-PEP8 | MAST2-DPR3 | EEDEYE | 30.7 [30.7 - 30.7] | 21.8 | - |
| MAST2-PEP9 | MAST2-DPR3 | DDDEYE | 29.4 [29.4 - 29.4] | 20.5 | - |
| MAST2-PEP10 | MAST2-DPR2 | IGEEDL | 32.2 [32.2 - 32.3] | 21.5 | Leu (P<sup>0</sup>) |
| MAST2-PEP11 | MAST2-DPR2 | HGEEDL | 31.6 [31.6 - 32.4] | 20.8 | Leu (P<sup>0</sup>) |
| MAST2-PEP12 | MAST2-DPR2 | QGEEDL | 31.4 [31.4 - 31.9] | 20.7 | Leu (P<sup>0</sup>) |
| MAST2-PEP13 | MAST2-DPR2 | VGEEDL | 31.2 [31.2 - 31.2] | 20.5 | Leu (P<sup>0</sup>) |
| MAST2-PEP14 | MAST2-DPR2 | MGEEDL | 31.0 [31.0 - 31.7] | 20.3 | Leu (P<sup>0</sup>) |
| MAST2-PEP15 | MAST2-DPR1 | EYGDMD | 30.3 [30.2 - 31.0] | 19.7 | Met (P<sup>-1</sup>) |

**Appendix Table 4:** Results of validation and control experiments with D-amino acid GyGlanvdessG, DPRV, and ST0929 control experiment scaffold bound to streptavidin. Peptide: the name of the D-peptide. Sequence: the sequence of the D-peptide. Log_10_ K* score: the predicted binding affinity using the K* algorithm (see Appendix A). Trp79 contacts in B barrel: boolean value stating if the D-peptide establishes hydrophobic interactions with streptavidin Trp79. # H-bonds in flexible loop: the number of hydrogen bonds established between the D-peptide and the flexible loop, comprised of streptavidin Ser45-Ser52.

| Peptide | Sequence | K* score (log<sub>10</sub>) | Trp79 contacts in β barrel | # H-bonds in flexible loop |
| --- | --- | --- | --- | --- |
| GyGlanvdessG | GLANVDESS | 32.2 | Yes | 1 |
| ST0929 (control) | GLANVDESS | 26.6 | Yes | 1 |
| DPRV | WWMIGDWND | 32.8 | Yes | 2 |

**Appendix Table 5.** Comparison of computational predictions versus experimental measurements for binding affinity of cRaf-RBD. The mutations column includes the wildtype and 41 test case mutations. The computational and experimental (%) columns represent the change in binding relative to wildtype (row 1). The rankings have a Spearman rank correlation coefficient of 0.81. This table is adapted from Lowegard et al.^23^.

| Mutation(s) | Experimental (%) | Computational (%) | Experimental Ranking | Computational Ranking |
| --- | --- | --- | --- | --- |
| Wild-Type | 100 | 100 | N/A | N/A |
| R89L | $1.3 \times 10^{-7}$ | $1.64 \times 10^{-10}$ | 1 | 3 |
| F61W/R67L/V69E/N71R/V88I/A85K | 0.23 | $2.2 \times 10^{-11}$ | 2 | 2 |
| K84A | 0.93 | $3.03 \times 10^{-5}$ | 3 | 4 |
| Q66A | 1.76 | 0.99 | 4 | 16 |
| A85D | 3 | 0.01 | 5 | 10 |
| R59A | 3.42 | $8.09 \times 10^{-4}$ | 6 | 7 |
| F61W/V69E/N71R/V88I | 4.64 | 0.03 | 7 | 12 |
| R67A | 6.19 | $1.78 \times 10^{-4}$ | 8 | 6 |
| K84L | 8.6 | $1.56 \times 10^{-4}$ | 9 | 5 |
| Q66K | 9 | 1.78 | 10 | 18 |
| T68A | 10 | 4.03 | 11 | 20 |
| V88D | 10 | 0.05 | 12 | 13 |
| T68K | 11 | $5.63 \times 10^{-17}$ | 13 | 1 |
| V69A | 13.68 | 5.08 | 14 | 22 |
| A85I | 18 | 8.64 | 15 | 24 |
| K65A | 18.57 | 1.04 | 16 | 17 |
| K65E | 19.4 | 0.71 | 17 | 15 |
| N64A | 21.31 | 5.77 | 18 | 23 |
| V69R | 29 | $1.4 \times 10^4$ | 19 | 34 |
| K87Q | 30 | 12.43 | 20 | 26 |
| K65M | 31.71 | 2.53 | 21 | 19 |
| N71E | 34 | 0.15 | 22 | 14 |
| F61W | 36.11 | 116.2 | 23 | 29 |
| F61W/R67L/N71R/V88I | 36.11 | 0.01 | 24 | 11 |
| V88I | 39.39 | 16.15 | 25 | 28 |
| R67L | 42 | $3.06 \times 10^{-3}$ | 26 | 9 |
| R59L | 43 | $2.51 \times 10^{-3}$ | 27 | 8 |
| K84R | 49 | 10.01 | 28 | 25 |
| N64D | 50 | 15.6 | 29 | 27 |
| F61W/N71R/V88I | 54.17 | $1.96 \times 10^4$ | 30 | 35 |
| K87R | 100 | 120.04 | 31 | 30 |
| F61W/N71R | 162.5 | $1.1 \times 10^5$ | 32 | 37 |
| V88K | 171 | 127.37 | 33 | 31 |
| V88H | 266 | 227.92 | 34 | 32 |
| A85R | 290 | $1.5 \times 10^7$ | 35 | 39 |
| N71R | 325 | $9.97 \times 10^4$ | 36 | 36 |
| N64K | 380 | 4.47 | 37 | 21 |
| V88R | 400 | $2.44 \times 10^3$ | 38 | 33 |
| A85K/V88R | 550 | $1.33 \times 10^7$ | 39 | 38 |
| A85K | 700 | $2.13 \times 10^7$ | 40 | 40 |
| N71R/A85K | 866.67 | $3.63 \times 10^9$ | 41 | 41 |

**Appendix Table 6.**
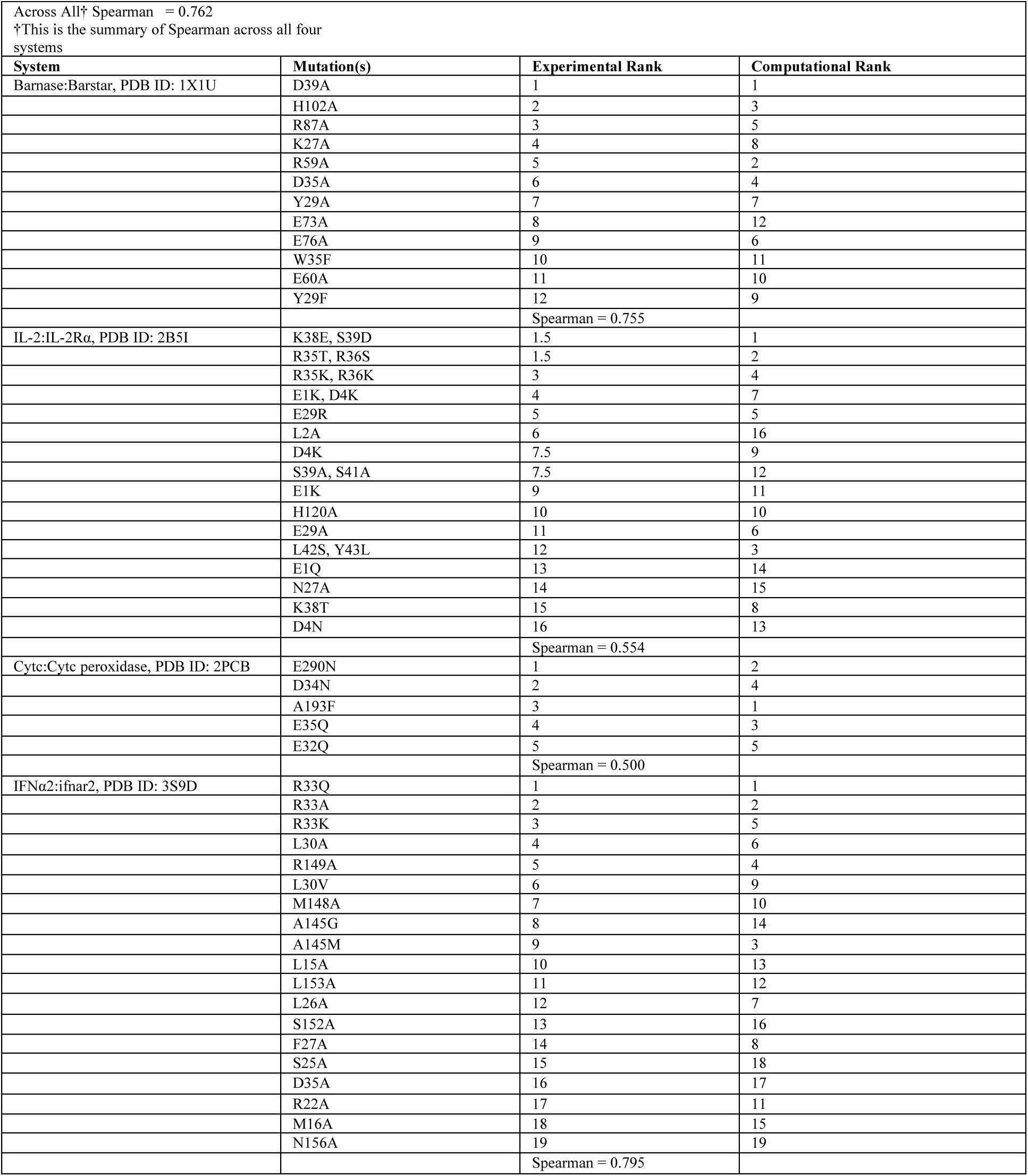
Comparing OSPREY binding affinity predictions to experimental measurements for four protein systems. The first column includes the system with relevant PDB ID, while the second describes the mutation. The computational and experimental rankings are provided in column three and four, respectively. Each system is divided by its individual Spearman rank correlation coefficient, with the first row providing the “across all” Spearman’s of 0.762. This table is adapted from Lowegard et al.^15^.

**Appendix Table 7.** Comparison of computed K* scores (log_10_) to experimentally-measured K_D_ for endogenous ligands and inhibitors of CAL and MAST2. Name: the name of the ligand. PDB ID: the reference ID for the structure in the Protein Data Bank^51^. Target: the protein the ligand binds and its function. Binding Affinity: the experimentally-determined binding affinity (inhibition constant or binding dissociation constant). K* Score (log_10_): the predicted binding affinity using OSPREY (see Appendix A for binding affinity calculation)^24^.

| Name | PDB ID | Target | Binding Affinity | K* Score (log<sub>10</sub>) |
| --- | --- | --- | --- | --- |
| CFTR C-terminus | 2lob | CAL (endogenous) | K<sub>i</sub> = 420 ± 80 μM<sup>25</sup> | 16.2 |
| kCAL01 | 6ov7 | CAL (inhibitor) | K<sub>i</sub> = 2.3 ± 0.2 μM<sup>21</sup> | 30.4 |
| PTEN6 | 2kyl | MAST2 (inhibitor) | K<sub>D</sub> = 1.99 μM | 28.8 |

### G. Appendix Figures

**Appendix Figure 1.**
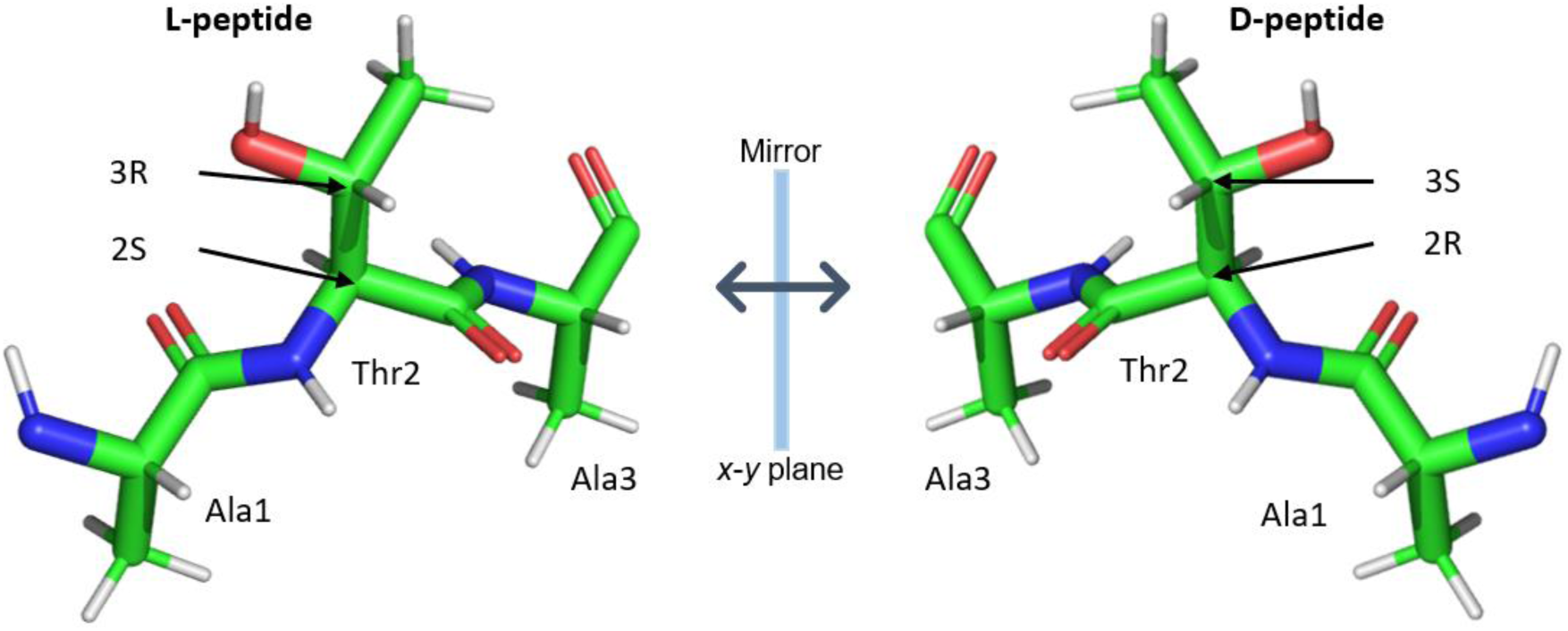
Reflection across a plane converts L-peptides to their D-counterparts, and vice-versa. Without loss of generality, here we reflect the L-tripeptide Ala-Thr-Ala (left) to its D-counterpart (right) across the *x*-*y* plane. Particularly noteworthy is that the reflection operation, implemented previously by Garton et al.^54^ for generation of a D-peptide database, correctly converts even L-amino acids with more than one stereocenter, namely threonine and isoleucine, to their D-counterparts. The second residue in the L-and D-peptides above is threonine. The absolute configuration of L-threonine’s two carbon stereocenters is (2*S*,3*R*). In contrast, D-threonine’s two carbon stereocenters have an absolute configuration of (2*R*,3*S*). As shown in the figure and described above, reflecting L-threonine, specifically (2*S*,3*R*)-2-amino-3-hydroxybutanoic acid, across the *x*-*y* plane obtains the correct enantiomer, namely D-threonine or (2*R*,3*S*)-2-amino-3-hydroxybutanoic acid. We have also confirmed that starting with L-isoleucine, specifically (2*S*,3*S*)-2-amino-3-methylpentanoic acid, and reflecting across the *x*-*y* plane obtains the correct enantiomer, namely D-isoleucine or (2*R*,3*R*)-2-amino-3-methylpentanoic acid.

**Appendix Figure 2:**
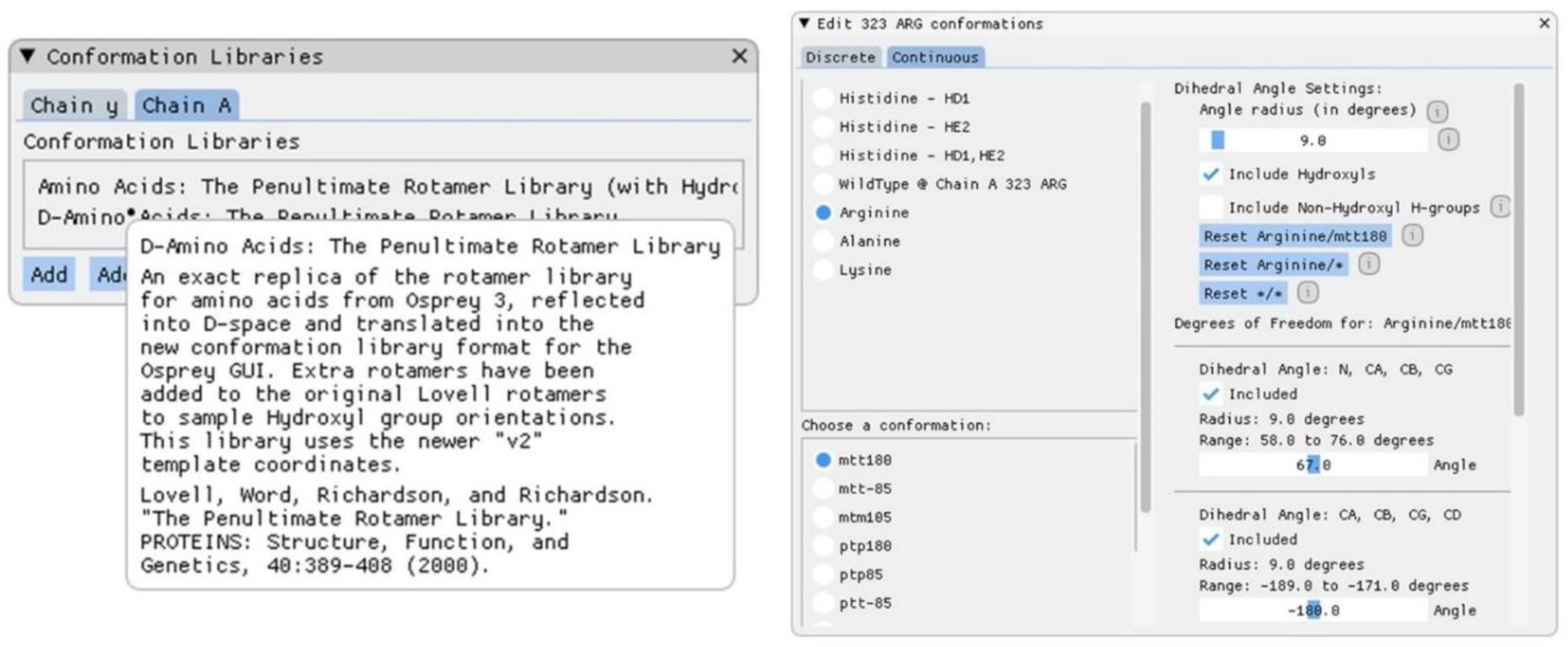
Screenshots of new OSPREY protein design specification options. OSPREY^1^ now allows protein designers to add their own conformation libraries and easily control rotamer selections and allowed movements. Left: DexDesign includes, in addition to the standard L-conformation library, a D-conformation library that is the mirror image of the L-library. A conformation library describes the standard connectivity templates, rotamers, and allowed movements, all of which can be further customized by the protein designer. A protein designer can specify multiple, distinct conformation libraries per chain. Right: New detailed control over side chain conformational flexibility. Each of the conformation library rotamers (e.g., tptm, pttm, etc.) can be included or excluded. The angle of voxel in which OSPREY continuously minimizes a rotamer^5^ can now be set, and each dihedral angle can be included or excluded from the continuous minimization. The customizations available to the protein designer with DexDesign unlocks the ability to explore new types of conformation spaces, such as the D-space.

**Appendix Figure 3:**
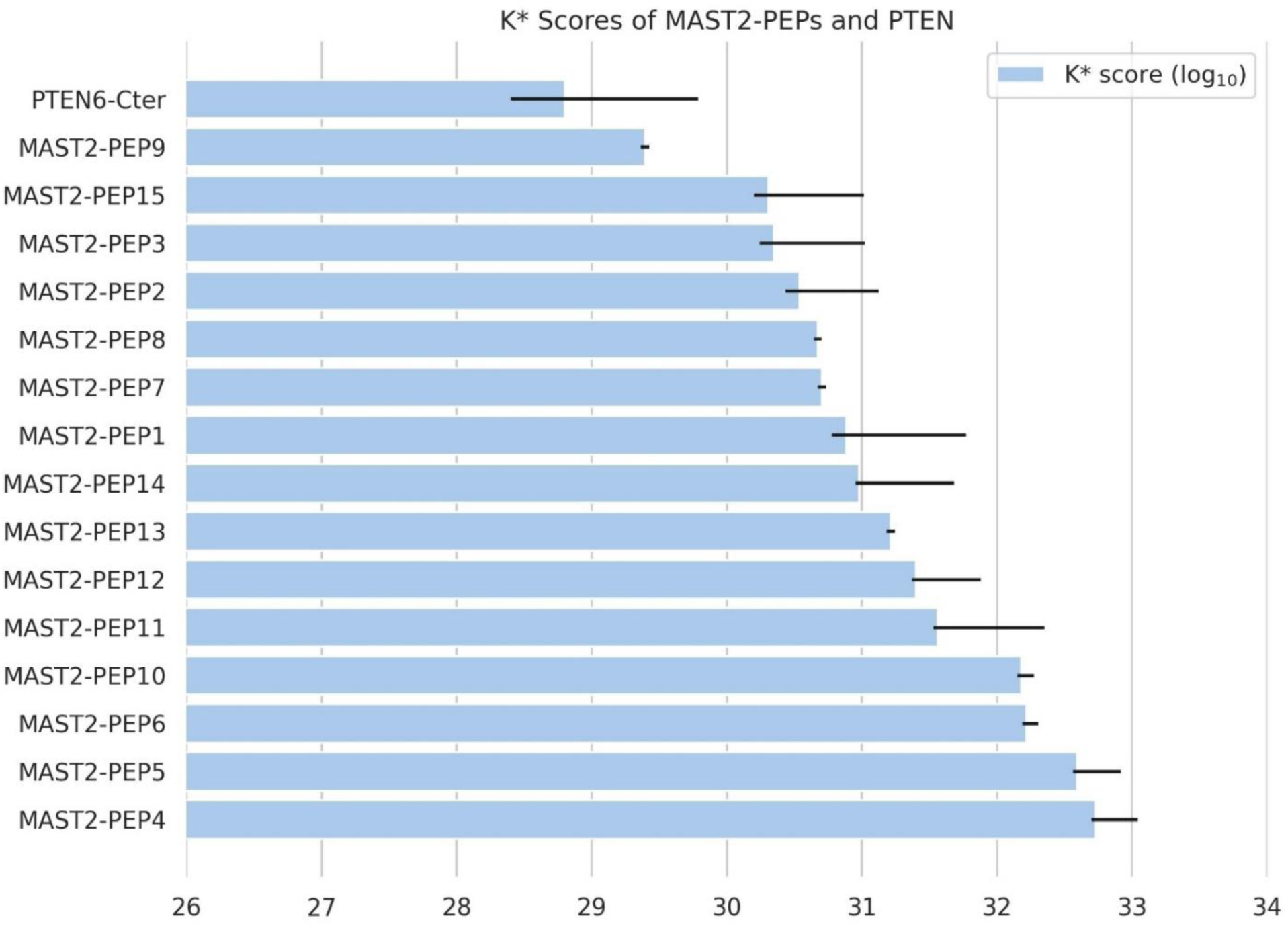
The DexDesign-generated D-peptides are predicted to bind to MAST2 tighter than PTEN. Blue bars show the OSPREY-predicted binding affinity of the MAST-PEPs and PTEN6-Cter (hereafter denoted PTEN6, the 6 C-terminal residues of MAST2’s endogenous ligand PTEN). We used the DexDesign algorithm (described in Section 2.1) and novel design techniques (described in Section 2.3.1) to generate 15 *de novo* D-peptides predicted by the K* algorithm^2,5^ to bind MAST2 tighter than PTEN6. Notably, PTEN binds MAST2 209-fold tighter than CFTR binds CALP (K_D_ = 1.9 ± 0.05 µM vs. 420 ± 80 µM)^25,55^, and binds MAST2 as tightly as the strongest known inhibitor of CALP, kCAL01 (K_D_ = 2.3 ± 0.2 µM)^21^, indicating a more challenging target of inhibition. OSPREY predicts PTEN6 to bind MAST2 with a log_10_ K* score (a provably accurate ε-approximations to *K*_a_, see Appendix A) of 28.8. Despite the more difficult target, all the MAST2-PEPs have higher log_10_ K* scores than PTEN6, ranging from 29.4 for MAST2-PEP9 to 32.7 for MAST2-PEP4, meaning the MAST-PEPs are predicted to outcompete PTEN6 and inhibit PTEN6:MAST2 binding. The best predicted inhibitor, MAST2-PEP4, has a ΔΔG of -1.1 kcal/mol, improving K_D_ 5-fold compared to the PTEN6 (see Appendix B). The K* scores of the MAST2-PEPs and empirical structures were determined using the K* algorithm^2,3^ in OSPREY^1^. Appendix A provides a definition of the K* algorithm and K* score. The error bars on the K* scores show the provable upper-and lower-bound of the K* approximation.

**Appendix Figure 4:**
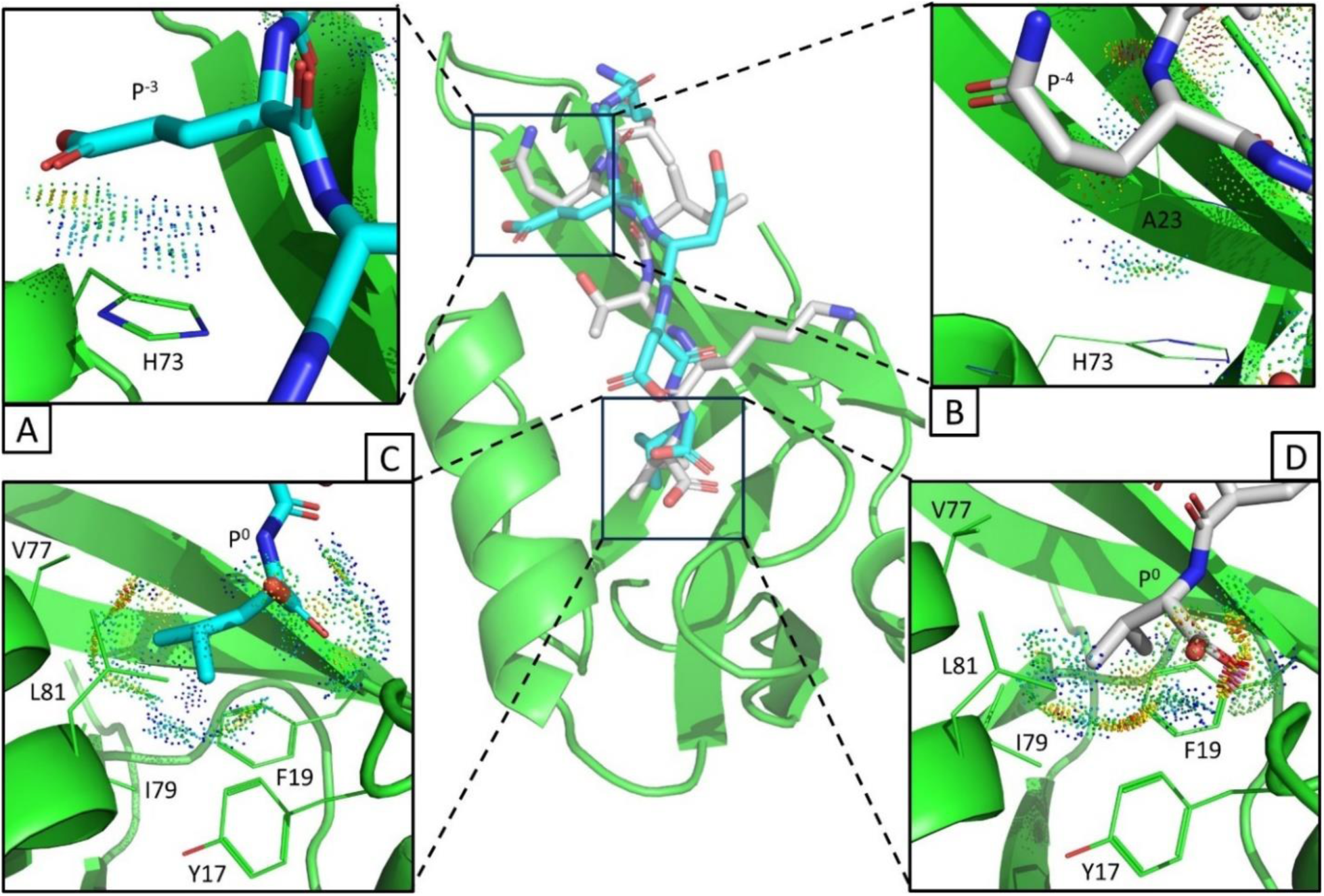
MAST2-PEP4 creates novel favorable interactions with MAST2 not found in PTEN. MAST2-PEP4 **(cyan sticks)** is the DexDesign-generated *de novo* D-peptide predicted to bind MAST2 **(green cartoon and lines)** with the tightest affinity. The K* algorithm^2,5^ in OSPREY^1^ predicts MAST2-PEP4 to bind MAST2 with a log_10_ K* score (a provably accurate ε-approximations to *K*_a_, see Appendix A) of 32.7, compared to 28.8 for PTEN6 (**gray sticks**, the 6 C-terminal residues of MAST2’s endogenous ligand PTEN). After normalization (see Appendix B), the Gibbs free energy change ΔG of the MAST2:MAST2-PEP4 complex is -8.8 kcal/mol, a -1.1 kcal/mol improvement over MAST2:PTEN6, resulting in a 5-fold improvement in K_D_. **Center:** the lowest-energy conformation from the OSPREY-predicted conformational ensemble of MAST2 **(green)** bound to MAST2-PTEN6 **(cyan)** and the lowest-energy model of PTEN6 **(grey)** from an empirical solution NMR ensemble of the MAST2:PTEN complex (PDB ID 2kyl)^28^. In comparison to the binding modes of, e.g., the CALP-PEPs to CALP (see Figure 5) which largely *recover* canonical PDZ-domain binding interactions, MAST2-PEP4 *restitutes* binding to MAST2 by exploiting novel geometric features of D-peptides not available to their L-counterparts (see Section 5). **A:** MAST2-PEP4’s P^-3^ glutamate makes favorable van der Waals contacts with MAST2’s His73 imidazole side chain, as indicated by predominantly green and blue MolProbity dots^29,56,57^. These favorable contacts are absent in the MAST2:PTEN6 complex. **B:** In contrast to **(A)**, PTEN6 cannot make some favorable interactions available to our D-peptides. For example, in MAST2:PTEN6, PTEN6’s P^-3^ isoleucine is oriented towards MAST2’s β2 strand (not shown), and the residue nearest to His73 is P^-4^ glutamine, whose amide fails to make van der Waals contacts with His73. **C:** MAST2-PEP4’s P^0^ leucine fills MAST2’s hydrophobic cavity^55^ formed by Tyr17, Phe19, Val77, Ile79, and Leu81. In contrast to PTEN6’s P^0^ valine **(D)**, MAST2-PEP4’s P^0^ leucine forms favorable interactions, as indicated by the green and blue MolProbity dots, with all 5 of the cavity’s hydrophobic residues. In addition, a rotation of the C-terminal carboxylate alleviates a steric clash (indicated by the red and pink MolProbity dots in **D**) with the carboxylate binding loop present in the MAST2:PTEN6 complex.

**Appendix Figure 5:**
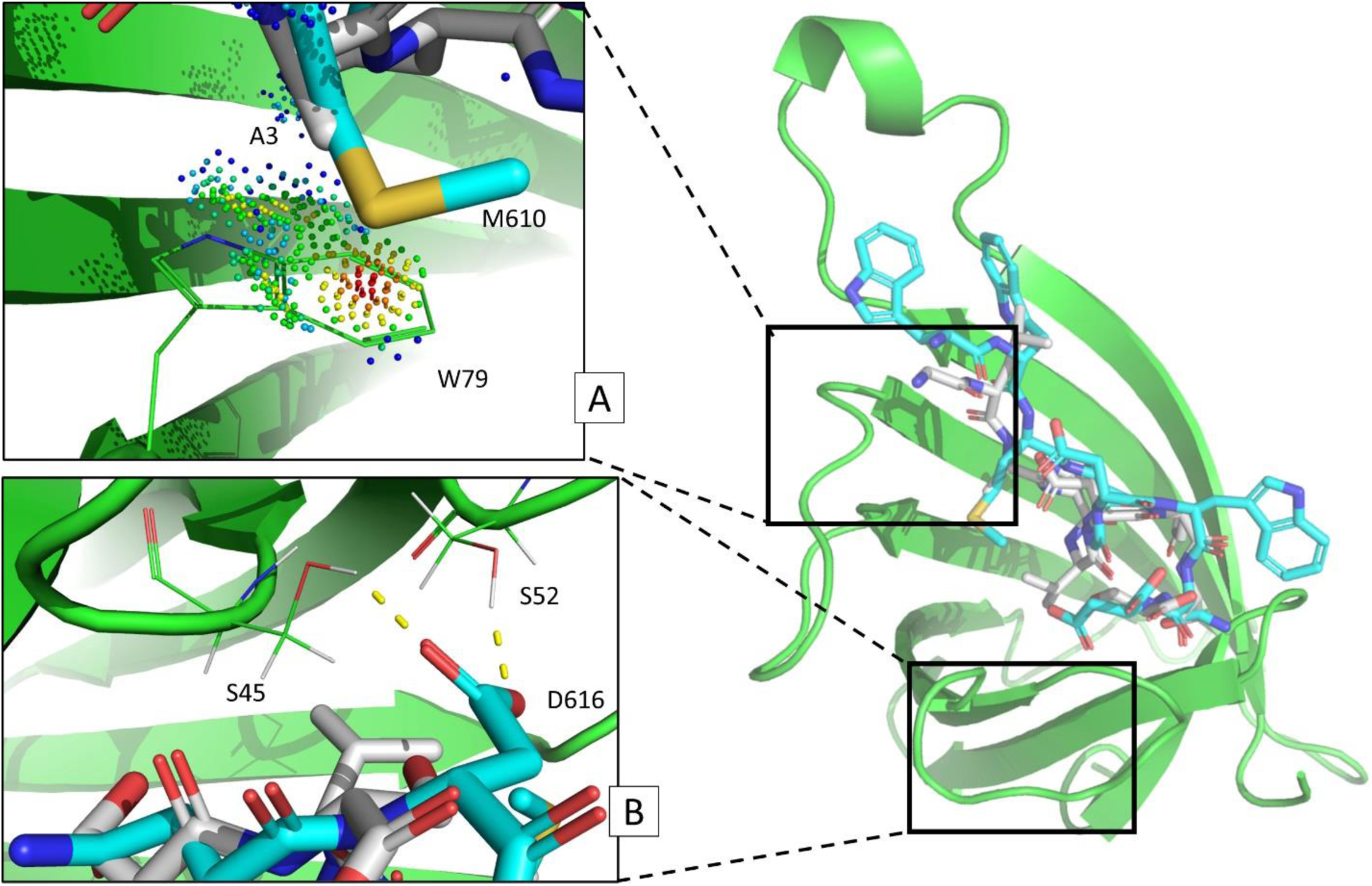
Geometric, chemical, and physical properties of DPRV that drive binding to streptavidin. **A:** Similar to GyGlanvdessG Ala3, DRPV Met610 displays favorable hydrophobic and van der Waals contacts with streptavidin Trp79. Streptavidin **(green cartoon and lines)** displays hydrophobic contributions through inward-facing tryptophan residues of the β barrel, which have been reported as important for ligand binding^37^. Favorable van der Waals interactions are shown as green and blue dots between DRPV **(cyan)** Met610 and streptavidin Trp79. GyGlanvdessG **(grey)** Ala3 also shares similar hydrophobic and van der Waals contacts with streptavidin Trp79. **B:** Differences in C-terminal orientation and interactions between D-peptide GyGlanvdessG:streptavidin and DPRV:streptavidin. Unlike GyGlanvdessG, DRPV’s aspartic acid C-terminus makes hydrogen bonds with both streptavidin residues Ser45 and Ser52. However, GyGlanvdessG’s C-terminal Ser9 fails to form a hydrogen bond with streptavidin Ser52. Residues number Ser45-Ser52 on streptavidin are known to be important for establishing binding of biotin^34^, so these contacts are likely evidence of high-affinity binding of DPRV.

**Appendix Figure 6:**
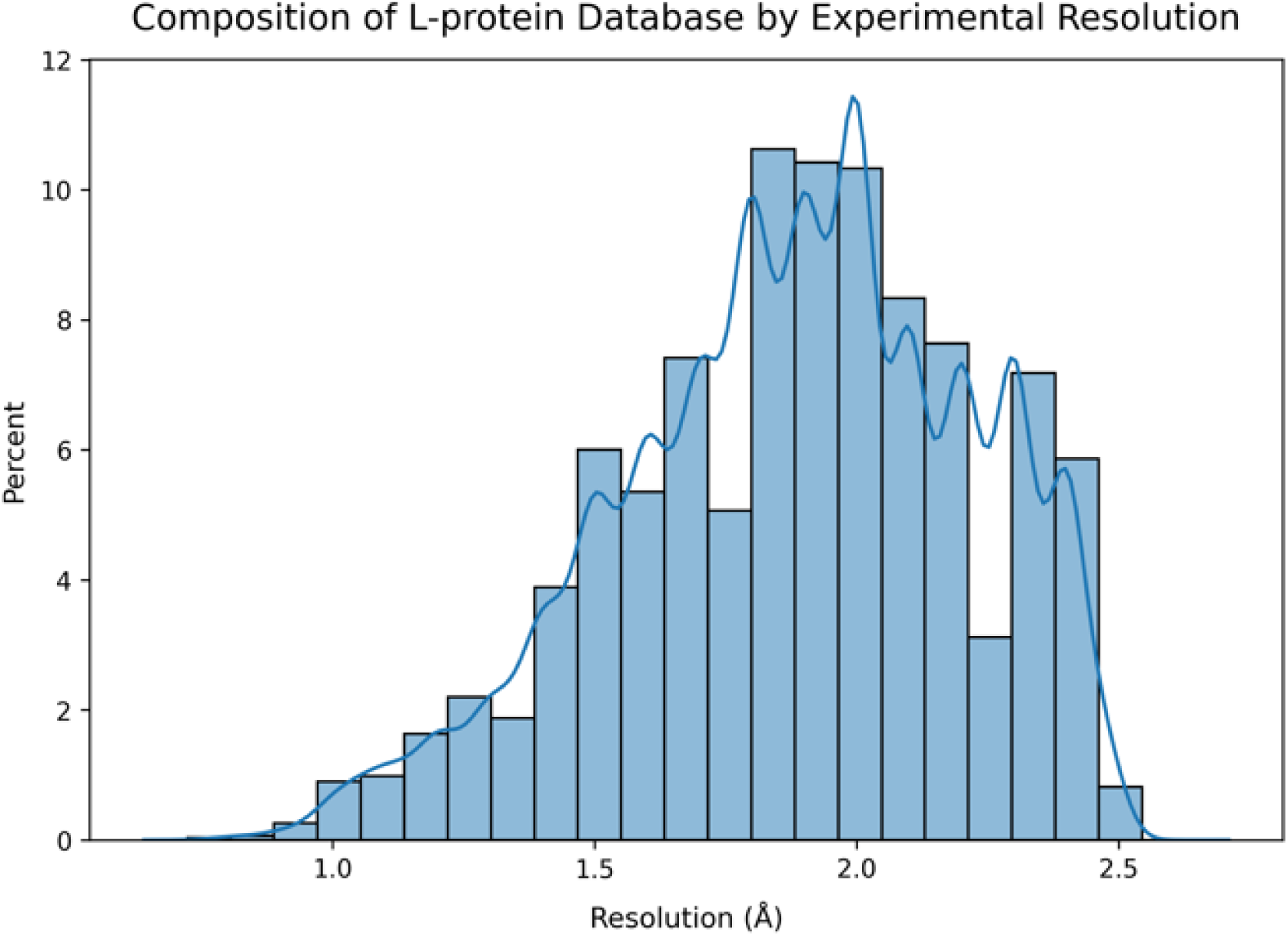
Composition of DexDesign’s MASTER search database by resolution. Step 3 of the DexDesign search algorithm executes a search over a protein structure database to identify L-protein segments with backbones similar (by RMSD) to the D-peptide query. For this, DexDesign uses MASTER^53^ to search over a protein structure database. We created a database of high-resolution L-protein structures by mining the RCSB PDB^51^ for crystallographically determined structures omitting DNA, RNA, and small molecules with a resolution of at most 2.5 Å. This resulted in a database containing 119,160 structures of varying resolutions. The histogram above shows the distribution of the resolution of the structures in the database we created. The mean, median, and model resolutions are 1.88 Å, 1.9 Å, 2.0 Å, respectively. The database was generated on 01/24/2023.

**Appendix Figure 7:**
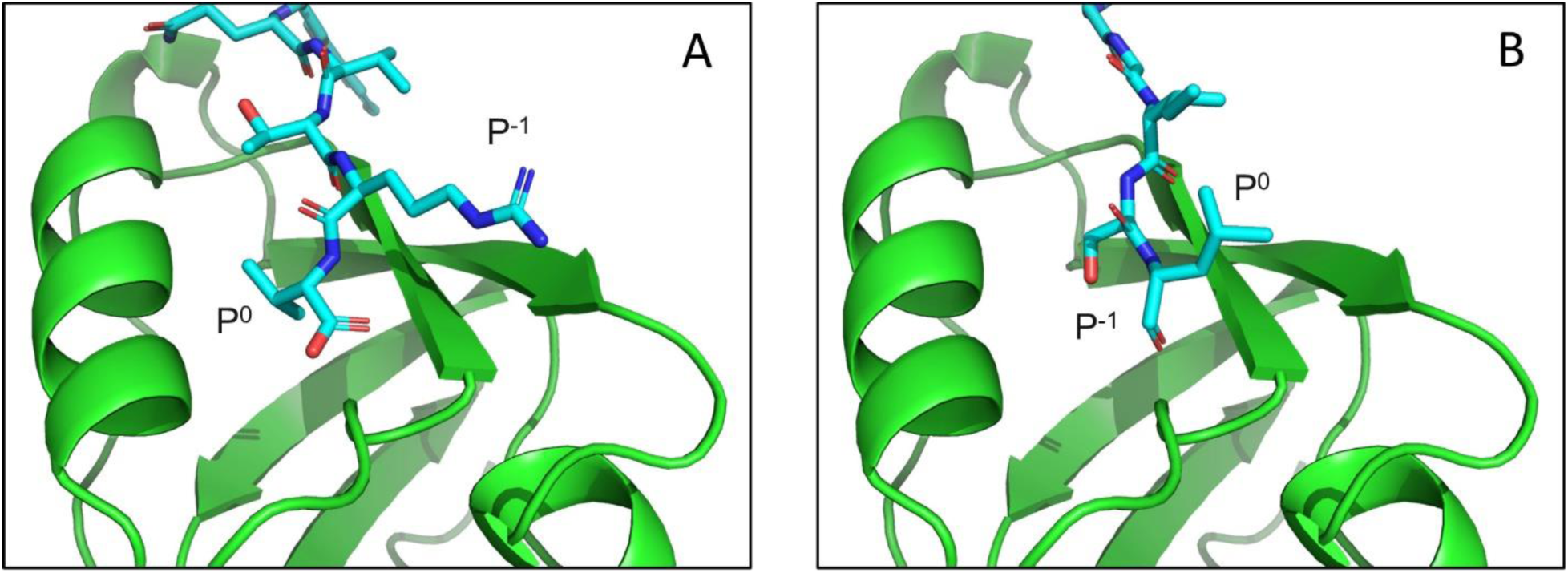
The sidechains of backbone-aligned L- and D- peptides point in opposite directions. **A:** sP^0^ and P^-1^ orientation for kCAL01 bound to CALP (PDB ID: 6ov7).^22^ The nonpolar P^0^ residue (Val) points towards CALP, filling the hydrophobic pocket. The charged P^-1^ residue (Arg) is oriented towards the CBL, where it forms three hydrogen bonds (not shown). **B:** P^0^ and P^-1^ orientation for CALP-DPR3 (D-form, PDB ID: 4m6r, residues A45-51, GGGISL) with CALP (L-form). In contrast to (A), the P^0^ residue (Leu) points toward the CBL. Furthermore, the P^-1^ residue (Ser) points toward the hydrophobic pocket. As expected, the residue positions are oriented in opposite directions between the L- and D- peptide, with an angle of 151° between kCAL01 and CALP-DPR3’s P^0^ residue and 155° between kCAL01 and CALP-DPR3’s P^-1^ residue. Therefore, redesign methods for CALP-DPR3 focus on filling the hydrophobic pocket with mutations and continuous flexibility at P^-1^ and establishing adequate hydrogen bonds between P^0^ and the CBL.

**Appendix Figure 8:**
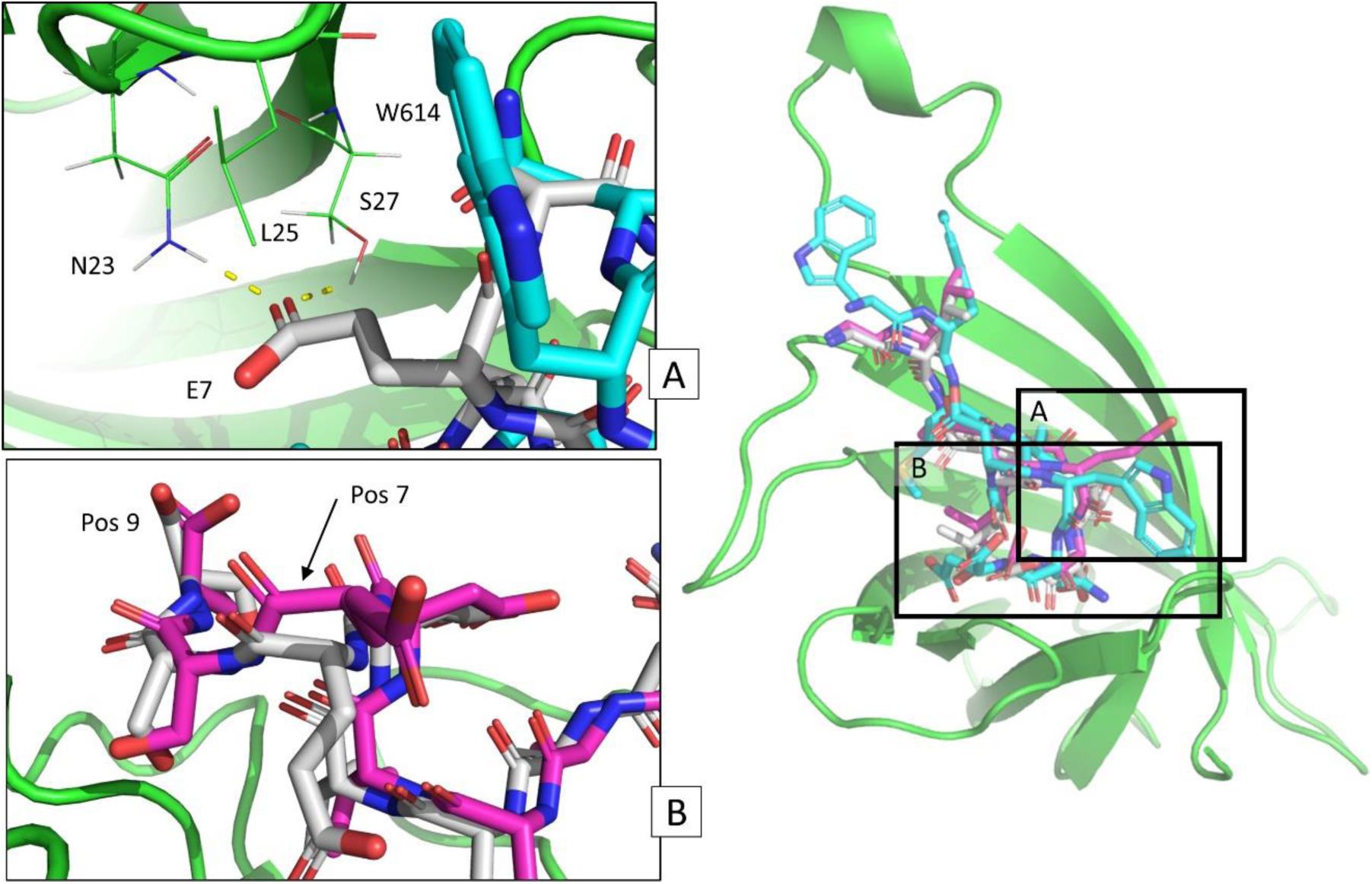
Backbone and hydrogen bonds at ligand residue positions 7 and 9 result in unique binding geometry and chemistry among D-amino acid GyGlanvdessG, DPRV, and ST0929 control experiment scaffold bound to streptavidin. **A:** Comparison of backbone orientation and hydrogen bonds between D-peptide GyGlanvdessG:streptavidin Glu7 and DPRV:streptavidin Trp614, which are both the 7^th^ residue from the N-terminus. The ST0929 control ligand is omitted for clarity. While GyGlanvdessG (grey) orients its Glu7 towards residues Asn23 and Ser27 in streptavidin (green cartoon and lines) to make two hydrogen bonds (yellow dashes), DPRV’s (cyan) Trp7 is unable to make contact with these residues. Were DPRV Trp614 to rotate towards these residues, it would encounter steric clashes with streptavidin residue Leu25. This is an example of irrecoverable ligand interactions due to different backbone geometries. B: Backbone geometry differences at residue positions 7 and 9 between GyGlanvdessG:streptavidin and the ST0929 control experiment scaffold bound to streptavidin. DPRV is omitted for clarity. The full backbone RMSD of the ST0929 control scaffold (magenta) to GyGlanvdessG (grey) is 0.48 Å. This seemingly small alignment difference yields a starting scaffold where 100% native sequence recovery results in a suboptimal binder, with the most notable backbone positions differences occurring at the C-terminus and 7^th^ residue from the N-terminus. The C-terminal oxygens point in different directions, with a distance of 1.8 Å between them. Further, the C_α_ carbons at the 7^th^ residue from the N-terminus are 1.2 Å apart. As described for DPRV in (A), residues at the C-terminus and 7^th^ residue position from the N-terminus exhibit either loss or generation of hydrogen bonds after application of the DexDesign protocol. Unlike GyGlanvdessG and similar to DPRV, the ST0929 control experiment scaffold is unable to make hydrogen bonds with streptavidin Ser27 (not shown). However, similar to GyGlanvdessG, it lacks a hydrogen bond in the flexible binding loop. Overall, this illustrates how differences in two key backbone positions can influence designability and predicted binding affinity.

**Appendix Figure 9.**
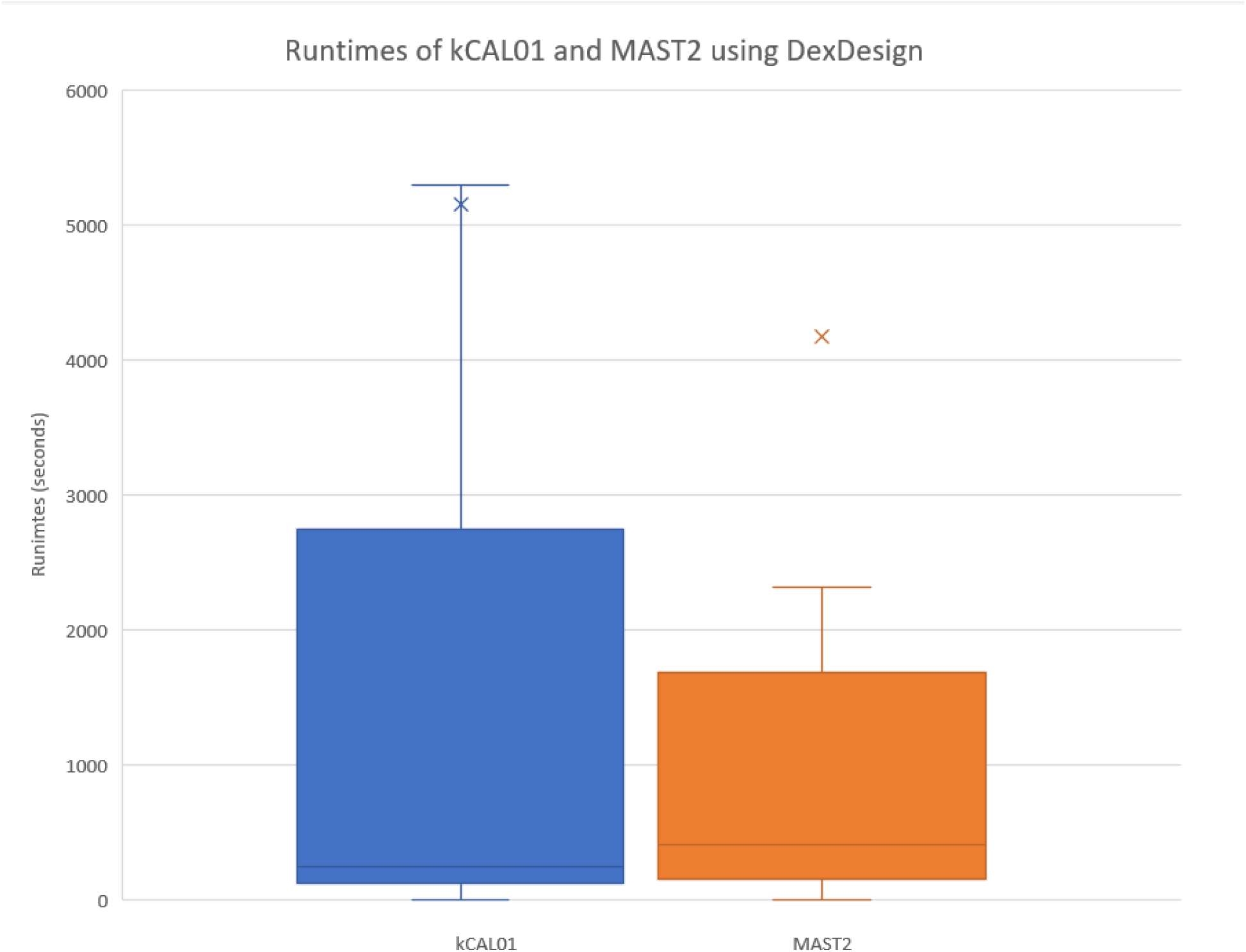
DexDesign Runtimes. Box-and-whisker plot showing the minimum, maximum, first quartile, and third quartile runtimes for DexDesign on CALP and MAST2. Each point represents the runtime, in seconds, for each design step in DexDesign. These steps include Minimum Flexible Set, Inverse Alanine Scanning, K* Optimizations, backbone minimization, and K*-based Mutational Scans. The runtimes for CALP and MAST2 ranged from 7 to 92,221 seconds and 3 to 113,490 seconds, respectively. Including all DexDesign steps averaged over both systems, the total runtime was 4,796 seconds, and the median was 326 seconds. For clarity, outliers are excluded in the figure. For example, the mutational scan for the 4^th^ residue from the N-terminus failed to finish in acceptable time for kCAL01-DPR3 and kCAL01-DPR5, which indicates a suboptimal location for mutation. These values are not shown in the graph. Finally, the fast median running times reported here are due to the fact that DexDesign implements exponential reductions in the search space from 20^6^ sequences to 3.8^3.4^ sequences. For more information, refer to Section 4 of the main text. These designs were run on a 24-core, 48-thread Intel Xeon processor with 4 Nvidia Titan V GPUs.

**Appendix Figure 10.**
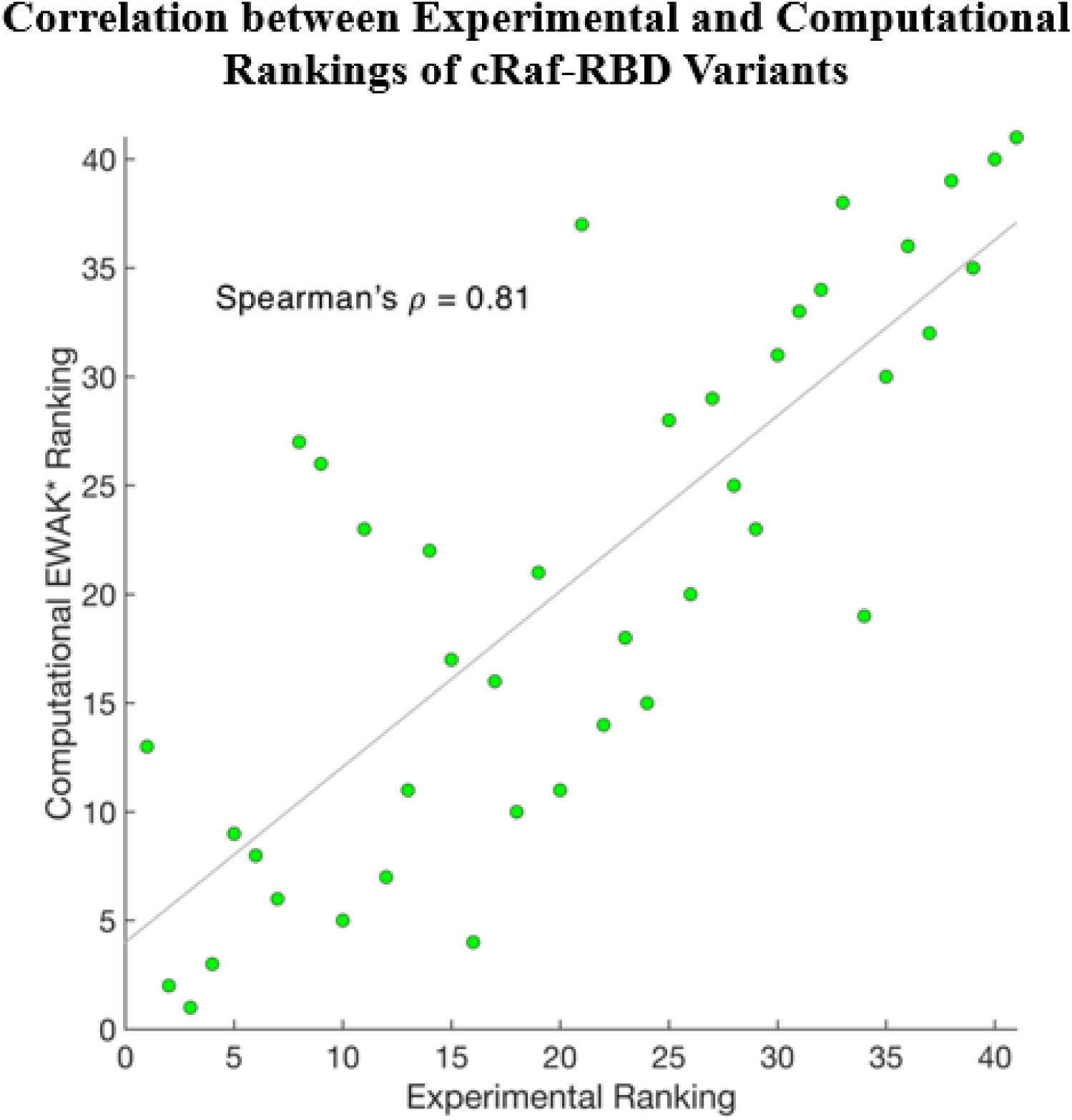
EWAK* (accelerated K*) ranking between computational predications of affinity and experimental measurements of affinity for 41 c-Raf-RBD variants binding to KRas. The horizontal axis includes experimental rankings, while the vertical axis includes computational (OSPREY) rankings as provided in Appendix Table 5. Each green dot represents a variant of c-Raf-RBD, and a least-squares fitting is provided via the solid grey line. The two rankings have a Spearman rank correlation coefficient of 0.81. This figure is adapted from Lowegard et al.^15^.

